# Mechanisms of AAV neutralization by human alpha-defensins

**DOI:** 10.1101/2024.09.25.614754

**Authors:** Jessica M. Porter, Kaitlin R. Hulce, Mackenzi S. Oswald, Kevin Busuttil, Shanan N. Emmanuel, Antonette Bennett, Robert McKenna, Jason G. Smith

## Abstract

Antiviral immunity compromises the efficacy of adeno-associated virus (AAV) vectors used for gene therapy. This is well understood for the adaptive immune response. However, innate immune effectors like alpha-defensin antimicrobial peptides also block AAV infection, although their mechanisms of action are unknown. To address this gap in knowledge, we investigated AAV2 and AAV6 neutralization by human neutrophil peptide 1 (HNP1), a myeloid alpha-defensin, and human defensin 5 (HD5), an enteric alpha-defensin. We found that both defensins bind to AAV2 and inhibit infection at low micromolar concentrations, similar to our prior studies of AAV6. While HD5 prevents AAV2 and AAV6 from binding to cells, HNP1 does not. However, AAV2 and AAV6 exposed to HD5 after binding to cells are still neutralized, indicating an additional block to infection. Accordingly, both HD5 and HNP1 inhibit externalization of the VP1 unique domain of both AAV2 and AAV6, which contains a phospholipase A_2_ enzyme required for endosome escape and nuclear localization signals required for nuclear entry. Consequently, both defensins prevent AAV from reaching the nucleus. Disruption of intracellular trafficking of the viral genome to the nucleus is reminiscent of how alpha-defensins neutralize other non-enveloped viruses, suggesting a common mechanism of inhibition. These results will inform the development of vectors capable of overcoming these hurdles to improve the efficiency of gene therapy.

**Author Summary:** AAVs are commonly used as gene therapy vectors due to their broad tropism and lack of disease association; however, host innate immune factors, such as human alpha-defensin antimicrobial peptides, can hinder gene delivery. Although it is becoming increasingly evident that human alpha-defensins can block infection by a wide range of nonenveloped viruses, including AAVs, their mechanism of action remains poorly understood. In this study, we describe for the first time how two types of abundant human alpha-defensins neutralize two AAV serotypes, AAV2 and AAV6. We found that one defensin prevents AAV binding to cells, the first step in infection, while both defensins block a critical later step in AAV entry. Our findings support the emerging idea that defensins use a common strategy to block infection by DNA viruses that replicate in the nucleus. Through understanding how innate immune effectors interact with and impede AAV infection, vectors can be developed to bypass these interventions and allow more efficient gene delivery.

## Introduction

Adeno-associated virus (AAV) is a non-enveloped single-stranded DNA virus that belongs to the genus *Dependoparvovirus* of the *Parvoviridae* family. Their small icosahedral T=1 capsid is comprised of three viral capsid proteins, VP1, VP2, and VP3, which assemble stochastically in an approximately 1:1:10 ratio [1, 2]. There are currently 13 known AAV serotypes which were isolated from both human and non-human primate species [3-5]. Due to their diverse tropism and inability to replicate independently, recombinant AAVs (rAAVs) are used as gene therapy vectors [6-8]. There are currently eight approved rAAV-based gene therapies, with more being tested in clinical trials [8]. However, poor transduction efficiency and interactions with the host immune system present hurdles that must be overcome to produce more efficient vectors [6, 7, 9, 10]. While the interactions of AAVs with components of the adaptive immune response such as anti-capsid neutralizing antibodies [11-13] and T cells [14, 15] have been previously investigated, the interactions of AAVs with innate immune effectors including antimicrobial peptides such as human α-defensins are vastly understudied.

Defensins are short, amphipathic, and cationic innate immune peptides that have been studied for their antimicrobial properties [16, 17]. They are broadly antimicrobial against bacteria and fungi as well as enveloped and non-enveloped viruses. Their structures consist primarily of β-sheets stabilized by three disulfide bonds [16, 18, 19]. Some defensins also form dimers, which are essential for their antiviral activity [20-23]. There are two types of defensins produced by humans, α- and β-defensins. Human α-defensins can be further categorized into two subtypes: enteric and myeloid. Enteric α-defensins, human defensin 5 and 6 (HD5 and HD6), are constitutively secreted by Paneth cells in the crypts of the small intestine at low mM concentrations and are also present in the genitourinary tract [24-27]. Myeloid α-defensins, human neutrophil peptides 1 to 4 (HNP1 to HNP4), are primarily expressed in neutrophils, where they are stored in cytoplasmic azurophilic granules that fuse with phagolysosomes following uptake of bacteria and viruses [16, 17, 19]. In addition, neutrophils release HNPs into the extracellular space during the formation of neutrophil extracellular traps [28]. Although plasma or serum concentrations of HNPs in healthy individuals are in the low nM range, they can be elevated in patients with liver disease, bacterial meningitis, and some cancers, for which they have been investigated as biomarkers [29-34]. Notably, plasma HNP1 concentrations as high as 49 µM have been measured in patients at the onset of sepsis [34]. HNPs are also found in fluids lining epithelial surfaces including tears, where HNP1 concentrations of 0.5 µM in normal tears are elevated 10-fold post-operatively or in some diseased conditions [35, 36], and in the epithelial lining fluid of the lungs, where HNP1 concentrations of 31 to 79 nM in healthy patients are elevated up to 2.5 mM in diseased states such as cystic fibrosis [37].

While both α- and β-defensins have been shown to have activity against bacteria and enveloped viruses, only α-defensins inhibit non-enveloped viral infection [18, 19]. Human adenovirus (HAdV), human papilloma virus (HPV), polyoma virus, and rotavirus (RV) have all been shown to be sensitive to neutralization by either HD5, HNP1, or both [38-45]. Previously, Virella-Lowell and colleagues found that AAV2 can be inhibited by HNPs present in bronchoalveolar fluid and by a mixture of purified HNP1 and HNP2 [37]. We recently showed that AAV1 and AAV6 are also neutralized by both HNP1 and HD5 [22]. In our previous studies with HAdV and HPV, we found that α-defensins perturb viral uncoating, therefore preventing productive infection [38, 40, 41, 46, 47]. A similar mechanism was determined for JC polyomavirus neutralization by HD5 [44]. These findings suggest the possibility of a common mechanism utilized by defensins to block non-enveloped viral infection. However, alternative mechanisms of neutralization have also been described for polyomaviruses [43], and the molecular basis for defensin-mediated inhibition of parvoviruses and rotaviruses has not been elucidated.

To begin to determine how α-defensins inhibit AAV, we focused on AAV2 and AAV6. We found that like the previously described effects of myeloid α-defensins HNP1 and HNP2 [37], the human enteric α-defensin HD5 also neutralizes AAV2. We then sought to determine the stage of the AAV cell entry pathway that is perturbed by each defensin. Although HD5 but not HNP1 could inhibit AAV2 and AAV6 from binding to cells, both defensins were able to neutralize AAV2 and AAV6 when added post-attachment, suggesting a common block at a downstream step. Consistent with this notion, we found that both defensins inhibit the externalization of the unique domain of VP1 (VP1u), a crucial step in AAV entry. This in turn prevents endosome escape and nuclear localization. Our findings support the concept of a common mechanism by which human α-defensins neutralize non-enveloped viruses by perturbing intracellular trafficking but also reveal a novel AAV-specific mechanism.

## Results

### Both HD5 and HNP1 bind to AAV2 and neutralize infection

Prior studies have shown that human enteric and myeloid α-defensins may differentially impact infection of non-enveloped viruses, including AAV1 [22, 38, 42, 44, 48]. Accordingly, we first sought to determine the effects of HD5 and HNP1 on AAV2 infection using the same physiologic defensin concentrations from our previous study of AAV1 and AAV6 [22]. When incubated with AAV2 on ice before the mixture is added to cells (pre-attachment), both HD5 [inhibitory concentration (IC_50_), 7.5 µM; 95% confidence interval (CI), 6.8 to 8.3 µM; Hill slope, -4.0] and HNP1 (IC_50_, 9.7 µM; 95% CI, <10.7 µM; Hill slope, -19.8) inhibit AAV2 infection with similar potency, and complete neutralization is achieved by 40 µM HD5 and 20 µM HNP1 (Fig 1A and 1B). We then quantified binding of each defensin to the AAV2 capsid by monitoring real-time kinetics of their interaction using surface plasmon resonance (SPR). Consistent with our studies of AAV1 and AAV6 [22], both defensins bind to the AAV2 capsid (Fig 1C and 1D). HNP1 [equilibrium dissociation constant (K_D_) = 0.32 ± 0.03 µM] bound with approximately 1.8-fold greater avidity than HD5 (K_D_ = 0.57 µM ± 0.06 µM), while the stoichiometry of HD5 (385 ± 5 molecules per capsid) was approximately 1.7-fold greater than HNP1 (227 ± 10 molecules per capsid). Thus, in contrast to our prior findings with AAV1 and AAV6 where both defensins bind to each serotype with similar avidity, HNP1 binds more tightly than HD5 to AAV2. However, as was true for AAV1 and AAV6, more HD5 than HNP1 binds to AAV2.

**Figure 1.**
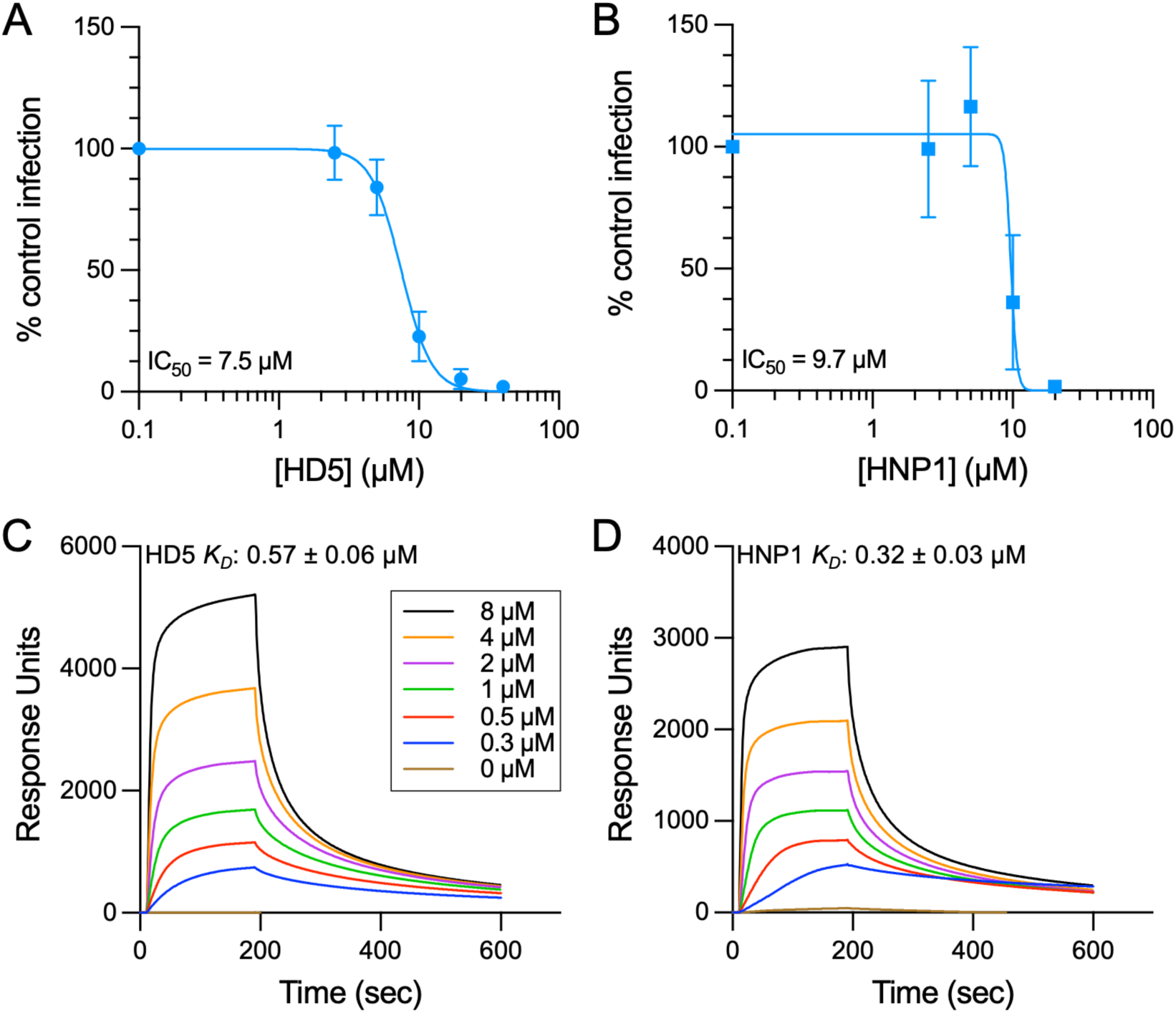
Both HD5 and HNP1 bind to AAV2 and neutralize infection. Infection of AAV2 in the presence of (A) HD5 or (B) HNP1. AAV2 was exposed to each defensin on ice prior to and during HeLa cell infection. Data are normalized to control infection in the absence of defensin and are the mean ± SD of a minimum of 3 independent experiments. Mean IC_50_ values are indicated. Both (C) HD5 and (D) HNP1 bind to AAV2. Background-corrected SPR sensorgrams are shown from a concentration series of defensin interacting with immobilized AAV2. Colors correspond to analyte (defensin) concentration, as indicated. Sensorgrams are representative of 2 independent replicates for each defensin. Mean *K_D_* values ± SD are indicated.

### HD5, but not HNP1, inhibits AAV2 binding to cells

To elucidate the mechanisms by which HD5 and HNP1 inhibit AAV2 infection, we first determined if either defensin could block AAV2 from binding to cells. We used qPCR to enumerate viral particles bound to cells in the cold in the presence or absence of increasing concentrations of defensin. HD5 (IC_50_, 9.2 µM; 95% CI, 6.7 to 14.1 µM; Hill slope, -3.7), but not HNP1, inhibited AAV2 cell binding (Fig 2A and B). Moreover, the IC_50_ of HD5 for binding (Fig 2A) and infection (Fig 1A) are equivalent (P = 0.94), suggesting that disrupted binding is the primary mechanism HD5 employs to block infection. We therefore reasoned that binding the virus to cells prior to defensin exposure might bypass inhibition of infection by HD5. However, contrary to our expectations, HD5 still neutralized infection when added post-attachment (IC_50_, 4.4 µM; 95% CI, 3.9 to 4.9 µM; Hill slope, -3.0) (Fig 2C). HNP1 also inhibited AAV2 under these conditions (IC_50_, 6.0 µM; 95% CI, 5.4 to 6.6 µM; Hill slope, -3.8), as expected (Fig 2D). Furthermore, as was true for AAV6 [22], both HD5 and HNP1 were more potent inhibitors of infection when added post-attachment rather than pre-attachment (HD5, P < 0.0001; HNP1, P = 0.0008). Thus, while inhibition of cell binding likely contributes to AAV2 neutralization by HD5, both defensins mediate a block of infection downstream of binding.

**Figure 2.**
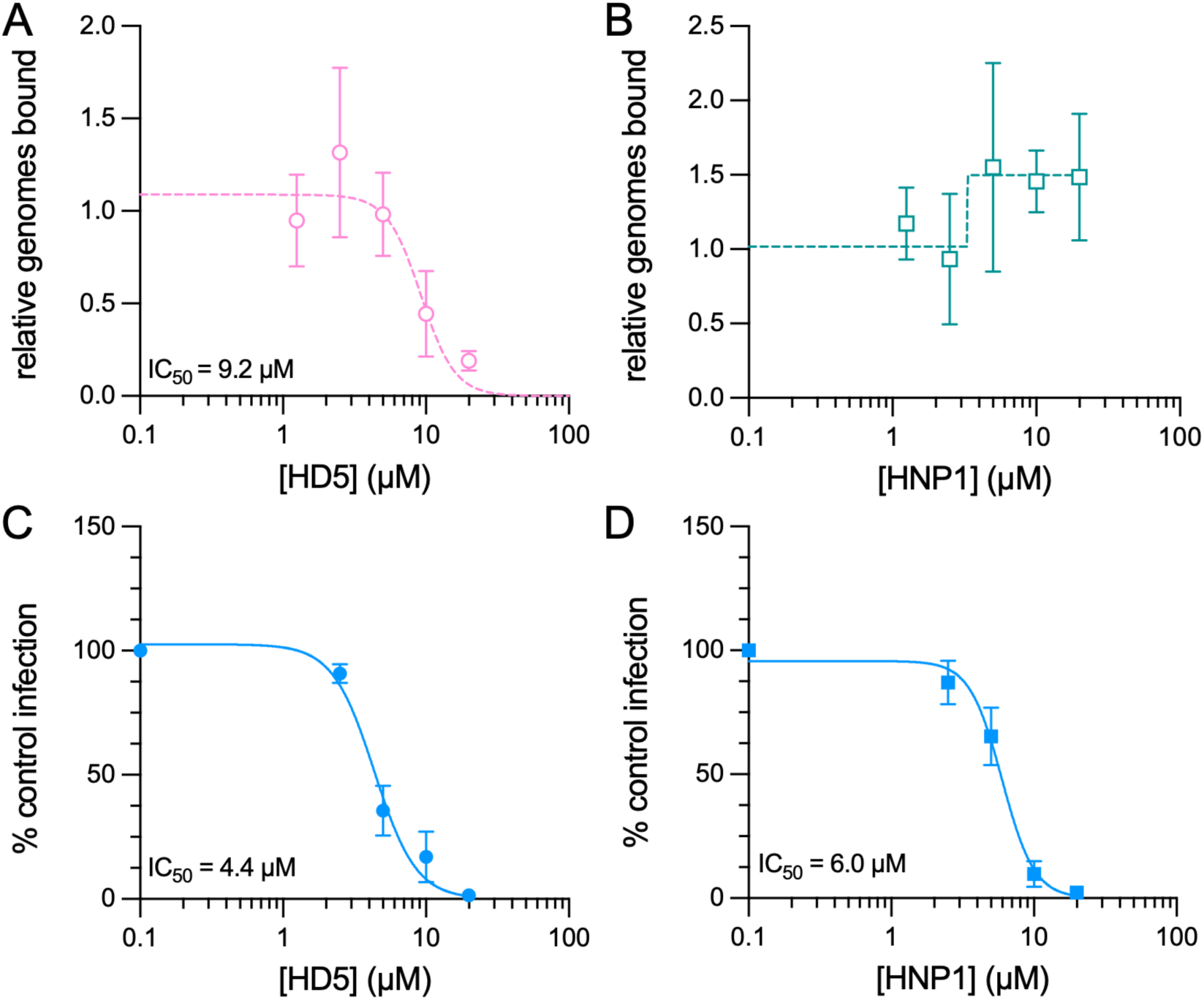
HD5, but not HNP1, inhibits AAV2 binding to cells, while both defensins impose a post-attachment block to infection. Effects of (A) HD5 and (B) HNP1 on AAV2 binding to HeLa cells. AAV2 was exposed to each defensin on ice prior to and during HeLa cell binding, which was then quantified by qPCR. Data are normalized to the amount of virus bound in the absence of defensin and are the mean ± SD of at least 3 independent experiments. The mean HD5 IC_50_ value is indicated. Both (C) HD5 and (D) HNP1 inhibit AAV2 infection when added after attachment of AAV2 to cells. Data are normalized to control infection in the absence of defensin and are the mean ± SD of a minimum of 3 independent experiments. Mean IC_50_ values are indicated.

### HD5 and HNP1 inhibit exposure of VP1u

A critical post-binding step in AAV entry is endosome escape mediated by the phospholipase A_2_ (PLA2) domain of VP1u. Before AAV enters the cell, VP1u is held within the capsid. However, conformational changes to the capsid occur within the endosome, resulting in externalization of VP1u [49-53]. To determine if this conformational change is disrupted by defensin interactions, we utilized antibodies [54] that bind either the externalized VP1u subunit (A1) or an epitope on the exterior of the VP3 domain (A20) in immunoprecipitation assays. HeLa cells infected with AAV2 in the presence or absence of an inhibitory concentration of HD5 or HNP1 were harvested 6 h post-infection (p.i.), and virus was precipitated from cleared lysate first by the A1 antibody then by the A20 antibody. Precipitated virus was visualized by immunoblot using the B1 antibody against VP1, VP2, and VP3 [54, 55]. Because VP3 is more abundant (approximately 10 VP3 to 1 VP1) and A1 binding to VP1u precipitates the entire capsid with which it is associated, we quantified the ratio of VP3 in the A1 blots to VP3 in the A20 blots across replicates. In the absence of defensin, both A1 and A20 bound AAV2 (Fig 3A-H). However, either 25 µM HD5 (Fig 3A and 3B) or 30 µM HNP1 (Fig 3C and 3D) was sufficient to block A1 binding, suggesting the absence of accessible VP1u. These defensin concentrations also completely inhibited infection under the conditions of this assay (Fig 3I-K). To exclude the possibility that the lack of A1 binding or accessibility was due to defensins competing with A1 for its epitope in VP1u, we infected cells in the absence of defensin, harvested at 5 h p.i. (for HD5) or 6 h p.i. (for HNP1), and then added HD5 or HNP1 to the cleared lysate during immunoprecipitation with A1 and A20. Under these conditions, both antibodies reacted with AAV2 (Fig 3E-H), indicating that the defensins are unable to compete with antibody binding. As an additional control, we utilized an HD5 analog (HD5abu) that is incapable of forming disulfide bonds due to the substitution of α-aminobutyric acid for the six cysteine residues in the linear sequence of HD5, rendering it non-functional as an antiviral (Fig 3I and 3K) [22]. HD5abu had no effect on A1 binding (Fig 3A and 3B), supporting the conclusion that the lack of A1 reactivity in the presence of HD5 or HNP1 is due to a specific effect of the defensins. From these experiments, we conclude that once the virus is internalized, both HD5 and HNP1 can prevent externalization of the VP1u subunit.

**Figure 3.**
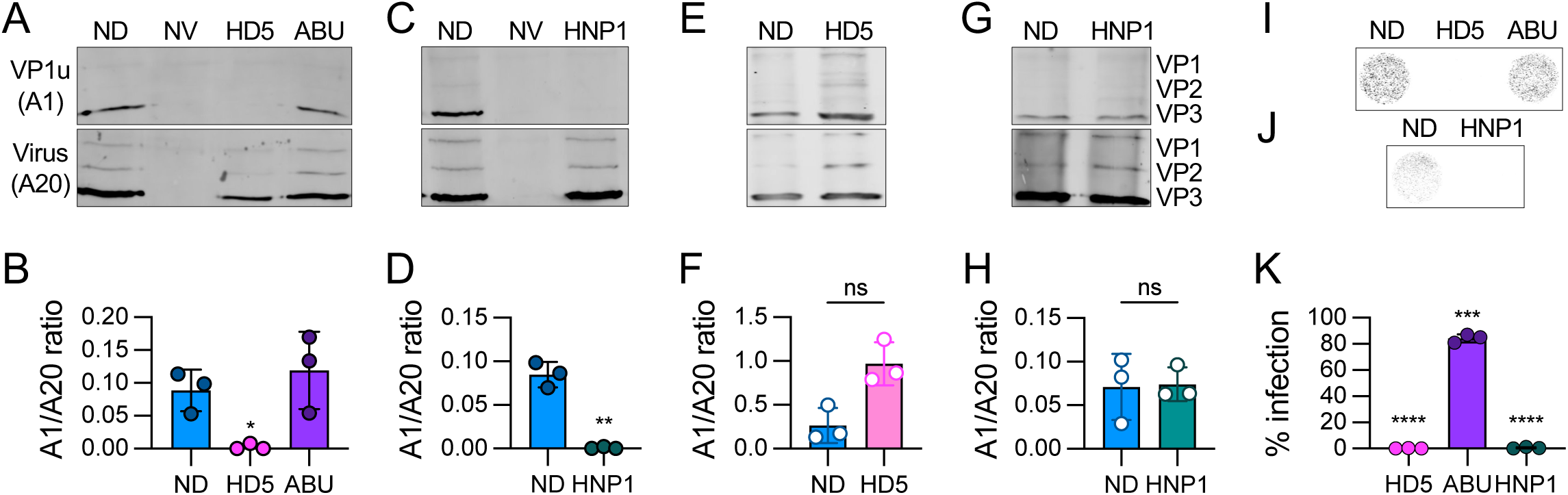
Both HD5 and HNP1 block AAV2 VP1u exposure. (A) Representative western blots of AAV2 that was sequentially immunoprecipitated at 6 h p.i. first with the A1 (VP1u-specific) antibody (top) then the A20 (intact capsid) antibody (bottom). Cells were infected in the presence of 25 µM HD5, 25 µM HD5abu (ABU), or no defensin (ND). Uninfected cells (no virus, NV) were included as a control. Blots were probed with the B1 antibody, and (B) the ratio of VP3 in the two blots was quantified. Positions of VP1, VP2, and VP3 for all blots are labeled in panel G. (C) Representative western blots and (D) quantification of similar experiments using 30 µM HNP1. (E) Representative western blots and (F) quantification of control experiments where no defensin was present at the time of infection, but 25 µM HD5 or no defensin (ND) was added to the cell lysate prior to immunoprecipitation. (G) Representative western blots and (H) quantification of similar experiments using 30 µM HNP1 post-lysis. (I and J) Representative images and (K) quantification of HeLa cells in 96-well plates infected with AAV2 in the presence or absence of the same concentrations of defensins that were used for immunoprecipitation. Images were obtained 48 h p.i. at a resolution of 50 μm, and grayscale intensity correlates with eGFP expression. Note that the infections were performed in parallel with the immunoprecipitation experiments but that the HNP1 infection experiments were done separately from those with HD5 and HD5abu. For B, D, F, H, and I, each point is a biological replicate (n = 3), and bars are the mean ± SD. Experiments in E and F used three times the culture size of the other panels, and infections were stopped 5 h p.i. instead of 6 h p.i.

To extend these findings to another serotype, we turned to AAV6, which is neutralized by both HNP1 and HD5 when added either pre- or post-attachment (Fig 4A and 4B and [22]). As was true for AAV2, HD5 but not HNP1 blocked AAV6 binding to cells (Fig 4A and 4B). However, while the correlation between inhibition of binding and infection by HD5 for AAV2 was 1:1, ∼4-fold more HD5 was required to block AAV6 binding (IC_50_, 41.6 µM; Hill slope, -0.9) than infection (IC_50_, 10.0 µM [22]). Because inhibition of cell binding by HD5 was incomplete, we performed additional control experiments with two known inhibitors of AAV6 binding and infection, the ADK1a neutralizing antibody [56, 57] and wheat germ agglutinin [58]. Both controls effectively blocked AAV6 binding, suggesting that the inability of HD5 to completely block AAV6 binding is not due to a limitation of the assay (Fig 4C). We then examined the effects of HD5 and HNP1 on VP1u exposure by AAV6. The A1 epitope is sufficiently conserved in AAV6; however, we used the ADK1a antibody in place of A20 to precipitate intact AAV6 capsids. We also performed immunoprecipitations 5 h p.i. and increased the scale of our experiments three-fold to maximize the A1 signal. We found that both defensins prevented AAV6 VP1u exposure, however, inhibition of both VP1u exposure (Fig 4D and 4F) and infection (Fig 4E and 4G) was less complete than we observed for AAV2. Taken together, we conclude that this mechanism of neutralization is conserved between at least two AAV serotypes.

**Figure 4.**
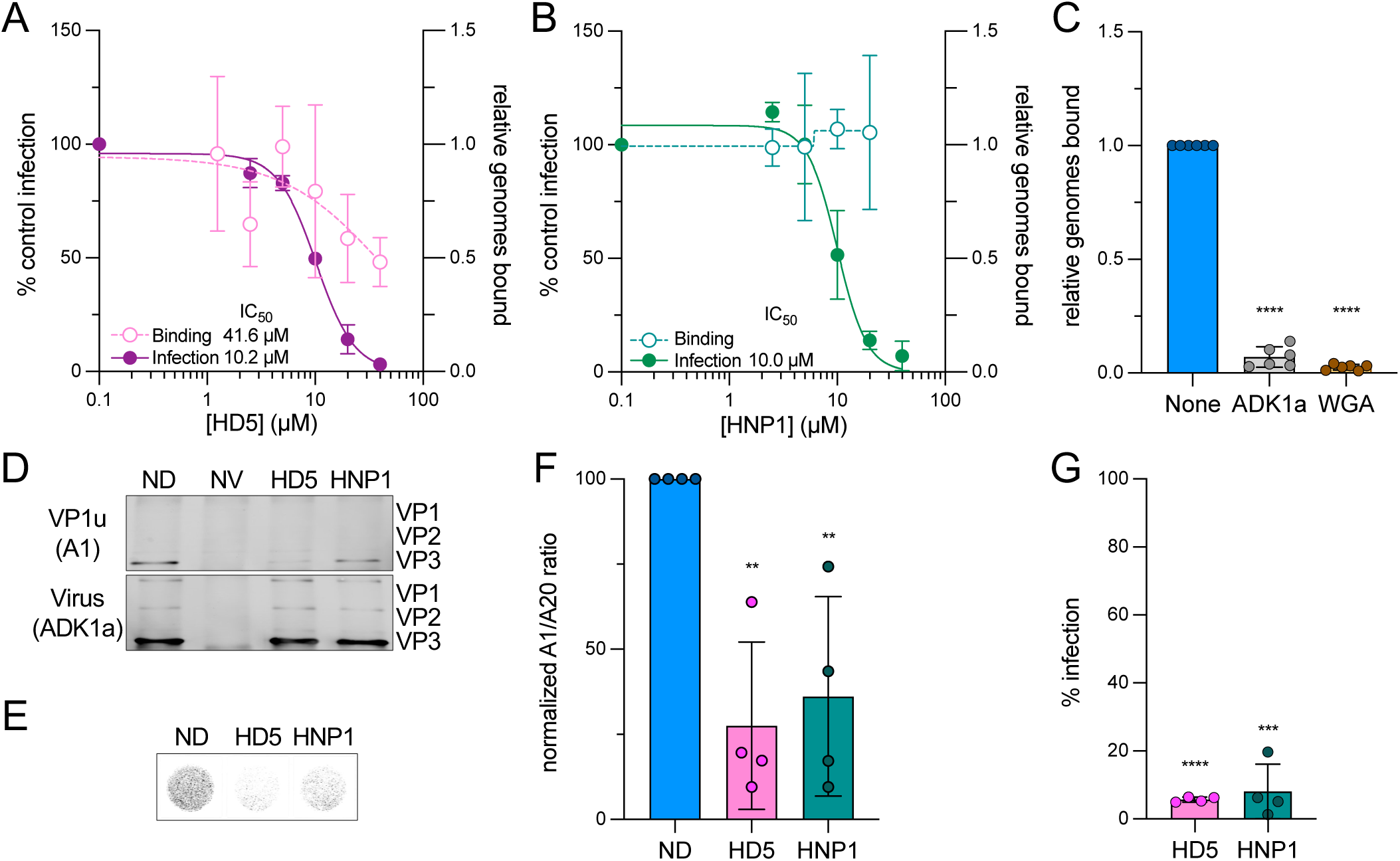
HD5, but not HNP1, inhibits AAV6 binding to cells, while both defensins block AAV6 VP1u exposure. Effects of (A) HD5 and (B) HNP1 on AAV6 binding to HeLa cells. AAV6 was exposed to each defensin on ice prior to and during HeLa cell binding, which was then quantified by qPCR. Data are normalized to the amount of virus bound in the absence of defensin and are the mean ± SD of at least 3 independent experiments. The effects of HD5 and HNP1 on AAV6 infection is reproduced from [22] for comparison. Data are normalized to control infection in the absence of defensin and are the mean ± SD of a minimum of 3 independent experiments. Mean IC_50_ values are indicated. (C) The neutralizing antibody ADK1a (1 µg/mL) and the lectin wheat germ agglutinin (WGA, 50 µg/mL) both block AAV6 binding to HeLa cells. (D) Representative western blots of AAV6 that was sequentially immunoprecipitated at 5 h p.i. first with the A1 (VP1u-specific) antibody (top) then the ADK1a (intact capsid) antibody (bottom). Cells were infected in the presence of 25 µM HD5, 30 µM HNP1, or no defensin (ND). Uninfected cells (no virus, NV) were included as a control. Blots were probed with the B1 antibody, and (F) the ratio of VP3 in the two blots was quantified then normalized to the ND control. (E) Representative images and (G) quantification of HeLa cells in 96-well plates infected with AAV6 in the presence or absence of the same concentrations of defensins that were used for immunoprecipitation. Images were obtained 48 h p.i. at a resolution of 50 μm, and grayscale intensity correlates with eGFP expression. For C, F, and G, each point is a biological replicate (n ≥ 3), and bars are the mean ± SD.

### Thermostability of the AAV2 capsid is not altered by either defensin

During these experiments, we noticed the appearance of a weak, virus-specific band in the A20-precipitated AAV2 samples that was only present in the absence of defensin (arrows, Fig 5A and 5B). Although it was not always observable in controls, likely due to the limit of detection of the assay, it was always absent or noticeably reduced in defensin-treated samples. The mobility of this band was faster than VP3, suggesting that it may be a proteolytic degradation product of one or more capsid proteins that retains the B1 epitope. We have previously described such degradation products of AAV2 capsids [59, 60]. The absence of this band in samples treated with either HD5 or HNP1 suggested that the defensins may be stabilizing the capsid and preventing proteolysis. Note that for AAV6, this region of the immunoblot is masked by strong signal from the ADK1a antibody heavy chain, making it difficult to assess if the same degradation product is present for this serotype. Thus, we focused follow-up studies on AAV2 to assess whether defensins alter capsid stability. To measure AAV2 capsid stability more broadly, we utilized differential scanning fluorimetry (DSF) in the presence or absence of either 25 µM HD5 or 30 µM HNP1 over a pH range from 4.0 to 7.4. This method uses the fluorescence of SYPRO Orange dye as a function of temperature to determine the melting temperature (T_m_) of the capsid. Although we observed a stabilizing effect of pH, with a maximal T_m_ of 78.2°C at pH 5.5 compared to 66.4°C at pH 7.4 and 74.2°C at pH 4.0, as observed previously [61], there was no effect of either defensin on thermostability (Fig 5C). These results suggest that defensin binding to the capsid might block specific conformational changes required for VP1u externalization or access of proteases to cleavage sites rather than creating an overall more stable capsid.

**Figure 5.**
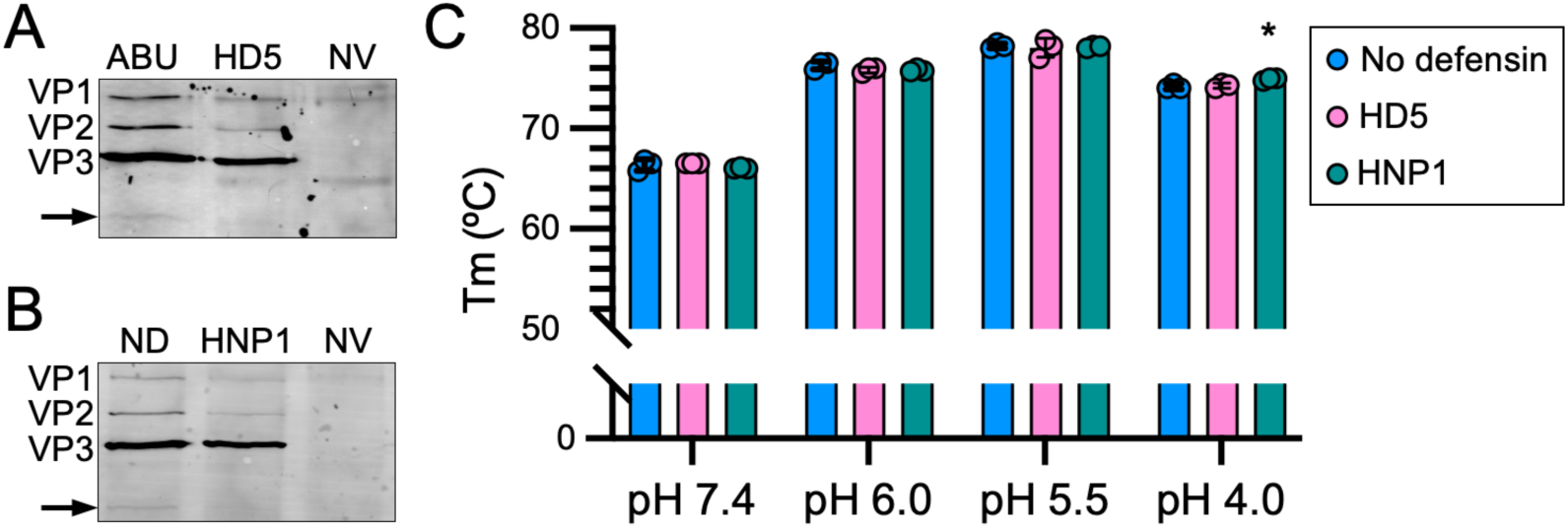
HD5 and HNP1 block proteolytic cleavage of the AAV2 capsid but do not alter capsid thermostability. Representative blots from additional replicates of the A20 immunoprecipitation experiments from Fig 3A and 3C probed with the B1 antibody. Arrows mark a B1-reactive proteolytic fragment of the capsid proteins that is detectable in (A) the presence of 25 µM HD5abu (ABU) but not 25 µM HD5 and (B) the absence (ND) but not presence of 30 µM HNP1. Uninfected cells (no virus, NV) were included as a control. (C) The melting temperature (T_m_) of AAV2 in the presence or absence of 25 µM HD5, 30 µM HNP1, or no defensin at the indicated pHs. Each point is a biological replicate (n = 3), and bars are the mean ± SD.

### AAV2 fails to enter the nucleus and accumulates perinuclearly in the presence of defensins

VP1u externalization serves at least two functions in AAV entry. The VP1u PLA2 domain disrupts the endosomal membrane, and VP1u contains nuclear localization signals that facilitate entry of the cytoplasmic capsid into the nucleus [50, 52, 62]. Therefore, we hypothesized that an inability of AAV2 to externalize VP1u in the presence of defensins would manifest as a reduction in nuclear localization of the capsid. We used confocal microscopy to visualize the trafficking of DyLight 488 (DL488)-labelled AAV2 in the presence and absence of a neutralizing concentration of either HD5 or HNP1. For these experiments, HD5 and HNP1 were added post-attachment due to the reduction in cell binding by HD5 when added to AAV2 pre-attachment. In the absence of defensin at 0 h p.i., DL488 AAV2 signal is diffuse with only coincidental colocalization with the nuclear marker (DAPI) in z-projected images (Fig 6A-C), consistent with the virus being bound to the outside of the cell. In contrast, at 24 h p.i., the viral signal is more punctate, and pronounced nuclear localization is observed. In the presence of either HD5 or HNP1 at inhibitory concentrations, AAV2 is punctate but exhibits markedly reduced nuclear localization. Rather, the virus accumulates perinuclearly, which we quantified by measuring the fraction of viral signal in each cell in a region spanning 5 pixels into and 50 pixels outside of the nucleus (Fig 6D). In untreated samples, the perinuclear fraction is smaller than the nuclear fraction; however, the ratio is reversed in HD5- or HNP1-treated samples (Fig 6E). Moreover, the absolute intensity of AAV2 at 24 h p.i. in defensin-treated samples is higher than in untreated samples (Fig 6F), which is consistent with reduced proteolytic degradation (Fig 5A and 5B). These data, in combination with the lack of A1 reactivity described above, support a model whereby both HD5 and HNP1 block VP1u externalization during entry, thereby impeding AAV2 from reaching the nucleus and neutralizing infection.

**Figure 6.**
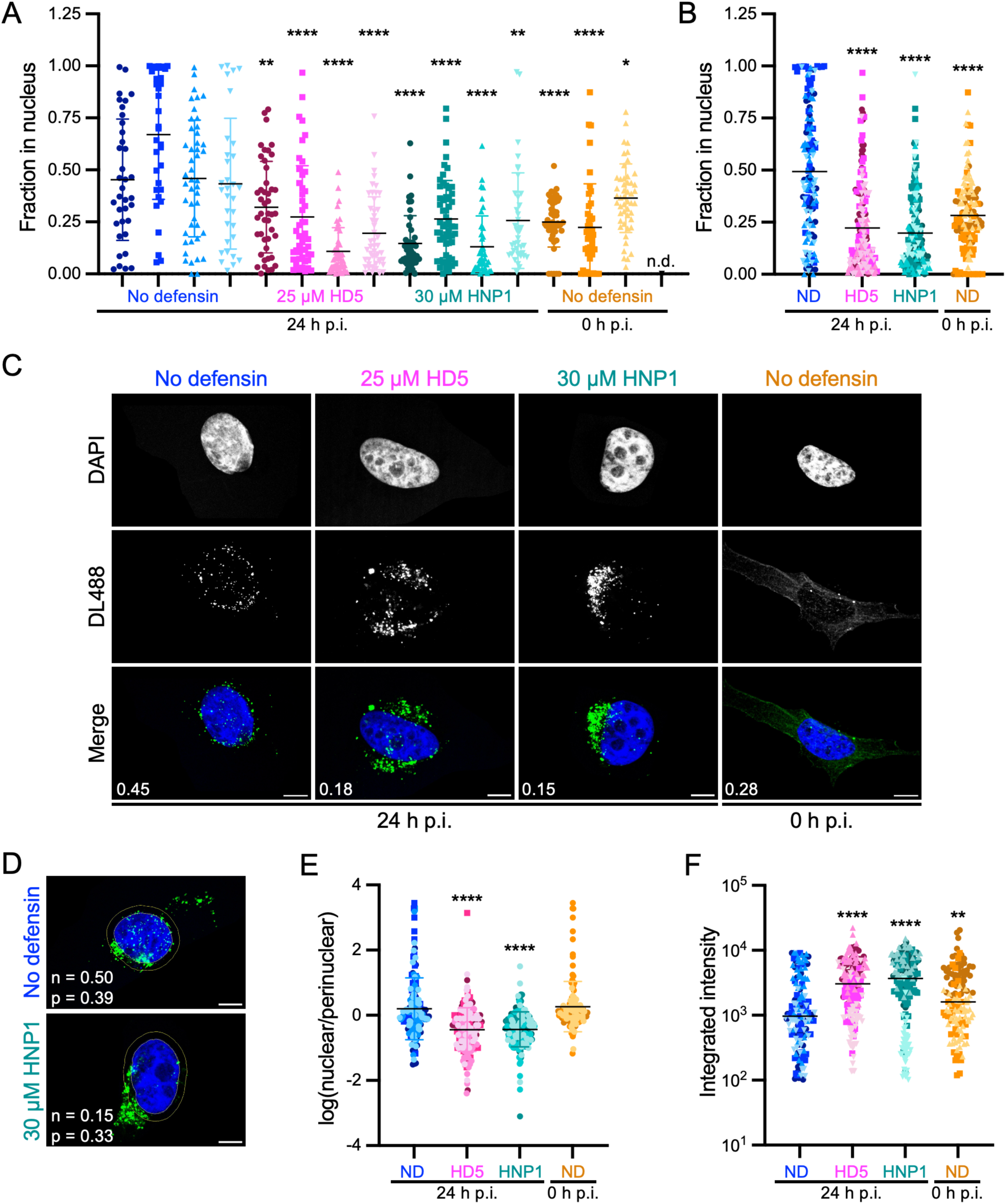
Both HD5 and HNP1 inhibit AAV2 nuclear localization. (A-C) The fraction of DyLight 488 (DL488)-labelled AAV2 in the nucleus of HeLa cells infected in the presence of 25 µM HD5 or 30 µM HNP1 was assessed 24 h p.i. by confocal microscopy and automated image analysis. Control cells infected in the absence of defensin (ND) were analyzed immediately after cold synchronization (0 h p.i.) or at 24 h p.i. (A) Scatter plots with a minimum of 35 cells per condition are shown. Each replicate is shown separately and indicated by both symbol (Replicate 1 = circles, 2 = squares, 3 = triangles, 4 = upside-down triangles) and color shade. n.d. = not done. (B) Replicates from A were pooled and re-analyzed in aggregate for each condition. C) Representative images of single cells from the experiments in A. Merged images include DAPI (blue) and AAV2-DL488 (green). The fraction of DL488 signal in the nucleus is indicated for each merged image. (D) Representative images of single cells from the experiments in A are shown to depict the perinuclear region, located between the concentric yellow rings, which was used for the analysis in E. The fraction of DL488 signal in the nucleus (n) and perinuclear region (p) is indicated for each image. (E) The integrated intensity of the DL488 signal in the nucleus was divided by that of the perinuclear region, and the log of this ratio is graphed. (F) The total cellular integrated intensity of the DL488 signal was quantified. In all graphs, horizontal bars indicate means. In B, E, and F, the shape and color of each data point corresponds to the replicates and conditions in A. In C and D, scale bars indicate 10 µm, the DL488 signal was individually adjusted for display purposes, and cells are from different replicates across conditions.

## Discussion

In this study, we have demonstrated that AAV2 infection is neutralized by both enteric and myeloid α-defensins with similar potency, extending our previous studies of AAV1 and AAV6 to an additional serotype [22]. In a normal infection, AAV binds to cells through interactions with attachment factors (e.g., heparan sulfate proteoglycans for AAV2 and proteoglycans with heparan sulfate or terminal sialic acids for AAV6) [63-65] and is endocytosed (Fig 7, left). In the endosome, the VP1u subunit, which was kept within the capsid prior to entry, is externalized [8, 49-52, 66]. This step is crucial for AAV entry, as the VP1u PLA2 domain mediates breakdown of the endosomal membrane, allowing virions to escape into the cytosol [53]. Host factors required for these steps are incompletely understood, but changes in pH and interactions with recently identified proteinaceous host factors including KIAA0319L (AAVR) and GPR108 play important roles [8, 66-70]. Once released from the endosome, AAV traffics along microtubules, and nuclear localization signals in VP1 and VP2 mediate entry of the virion into the nucleus [52, 62, 71-73]. The virion then uncoats in the nucleolus, releasing its genome to begin transcription and replication [74, 75]. In the presence of defensins, two crucial steps in this entry pathway are perturbed: cell binding and VP1u exposure (Fig 7, right). HD5 but not HNP1 perturbs cell binding, if encountered by AAV2 or AAV6 prior to cell attachment. However, if AAV2 or AAV6 succeeds in binding to the cell and becomes internalized, HD5 and HNP1 both prevent the externalization of the VP1u subunit downstream of cell binding, thereby trapping the virus in the endosome and blocking cellular trafficking and nuclear entry.

**Figure 7.**
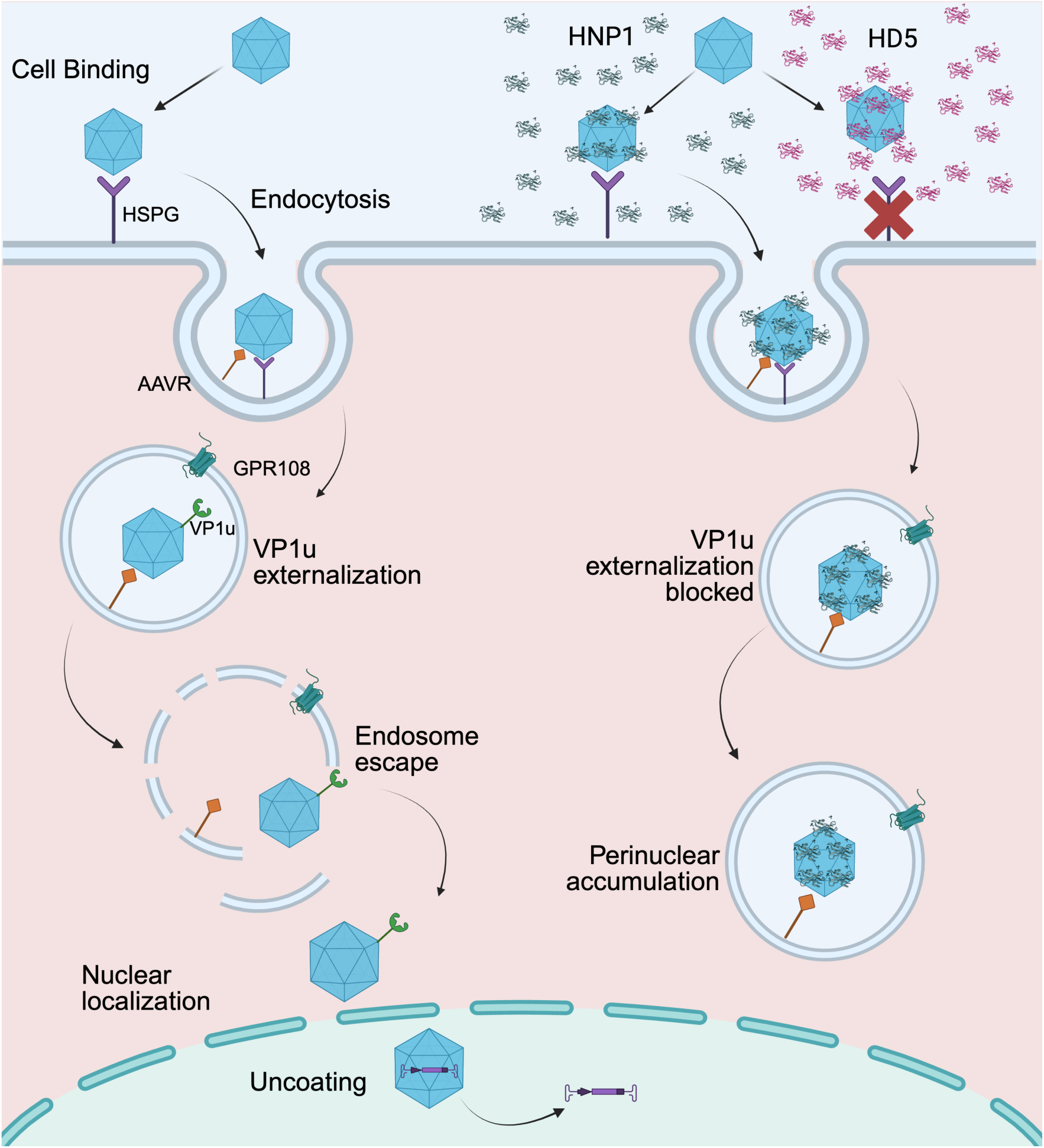
Model of AAV neutralization by defensins. Model of AAV infection in the absence (left) and presence (right) of neutralizing concentrations of HD5 or HNP1. Host factors including heparan sulfate proteoglycans (HSPG), the AAV receptor (AAVR), and GPR108 are depicted. The red “x” indicates the block to cell binding upon pre-attachment exposure of AAV2 or AAV6 to HD5. HNP1 is used to illustrate the effects of the intracellular defensin block, which is common to HD5 and HNP1. Figure created with BioRender.com.

How defensins interact with the capsid to block VP1u externalization remains unknown. A reasonable expectation is that both defensins must remain bound to the capsid at endosomal pHs, which are required to stimulate conformational changes required for VP1u externalization in the absence of defensin. However, it is possible that as the pH environment of the capsid changes during endocytosis, defensin binding is altered but not completely abrogated. There are at least two models by which defensin binding may prevent VP1u externalization: 1) altering the local chemical environment of pH-sensitive residues that form a “molecular switch” to trigger VP1u externalization [59], thereby impeding their protonation or 2) blocking the conformational changes that allow VP1u to be externalized through local stabilization or steric hinderance. These mechanisms are not mutually exclusive. One model is that VP1u is externalized through channels at the 5-fold axes that are too small to accommodate folded VP1u prior to conformational changes induced by the endosomal environment [76, 77]. In addition to localized changes at the 5-fold vertices, more extensive rearrangements involving other areas of the capsid including the 2-fold axis have been postulated [76, 78]. Our data suggest that defensins do not increase the overall thermostability of the AAV2 capsid, arguing against a model where the entire capsid is stabilized against host-induced conformational changes upon defensin binding. However, steric hinderance at the 5-fold axis pore may prevent either the necessary local conformational changes from occurring or block VP1u from threading through the pore. A similar model has been postulated for the anti-AAV8 and -AAV9 antibody HL2372 and the POROS CaptureSelect affinity ligand CSAL9, both of which bind to the 5-fold axis and neutralize infection [79-81]. In our prior studies of differences in defensin sensitivity of AAV1 and AAV6, the crucial residue was located at position 531 located on the arm of the 3-fold protrusion extending along the wall of the 2-fold axis [22]. A positively charged residue at this location (lysine) was associated with more uniform sensitivity to defensin neutralization. In contrast, viruses bearing a negatively charged residue (glutamic acid) had variable phenotypes dependent upon when defensin was added relative to cell binding. Therefore, we hypothesized that amino acid 531 is not directly involved in binding to cationic defensins. Rather, the nature of this residue may dictate surface charge at other parts of the capsid to modulate defensin interactions. Definitive mechanistic insight awaits structural information on the defensin-AAV complexes.

An additional hint to the location of defensin interactions with AAV may be obtained by further analysis of capsid protein proteolysis that appears to be blocked by both HD5 and HNP1 binding. Studies of purified virus have shown that AAVs possess intrinsic autoproteolytic activity that is activated at endolysosomal pH leading to degradation of VP1 and VP3 (and possibly VP2) [59] in addition to protease activity in the VP1u domain that is capable of cleaving external substrates (e.g., casein) [59, 82]. Thus, the absence of capsid degradation in the presence of α-defensins could constitute additional evidence that VP1u is not externalized and able to cleave the capsid. Alternatively, it could reflect direct inhibition of auto-proteolytic activity elsewhere in the capsid or capsid cleavage by cellular proteases during infection. In this regard, HD5 and HNP1 could block either the protease active site or the target cleavage site. To our knowledge, there have been no studies of inhibitors or mutants that have elucidated the role of capsid cleavage in AAV infection, although cleavage of the capsid proteins of autonomous parvoviruses by autoproteolysis or cellular proteases is an important step in maturation or entry [83-85]. Thus, further study of the inhibition of capsid cleavage may provide insight into where defensins bind to the capsid, but it is unclear if blocking proteolysis contributes to the defensin-mediated neutralization mechanism.

Although both HD5 and HNP1 block VP1u externalization, their interactions with the capsid differ. First, only HD5 disrupts AAV2, and less potently AAV6, from binding to cells. This mode of neutralization by α-defensins is uncommon, having only been described previously for BK polyomavirus, where HD5 aggregates the virus and reduces cell binding [43]. In contrast, cell binding of many viruses like HAdVs, mouse AdV, and HIV-1 is promoted by α-defensins [39, 46, 86-88]. Second, HNP1 binds AAV2 more tightly than HD5 does, but there is less HNP1 than HD5 bound to the capsid at equilibrium. These differences are not surprising given that the two defensins, although structurally similar, vary in primary sequence, net charge, and charge distribution. Unlike antibodies, defensins do not bind to a specific epitope. Rather, selective binding to favorable capsid regions may initially occur over a range of affinities. Primary binding may then promote additional defensin-defensin and defensin-capsid interactions. HD5, with weaker but more abundant binding, may sterically occlude heparan sulfate proteoglycan (HSPG) binding sites, inhibiting cell binding. HNP1 instead may bind preferentially to less abundant, higher affinity sites away from the HSPG binding site, allowing cell binding to occur. The importance of stoichiometry in blocking receptor binding is underscored by the phenotype of AAV6, where both defensins bind with similar affinity [22]. However, HD5 is not only more abundant on the capsid but also capable of blocking cell binding despite the ability of AAV6 to utilize either HSPG or terminal sialic acid for attachment. Thus, it is reasonable to postulate that the binding sites utilized by each defensin may not entirely overlap. These findings are reminiscent of HPV interactions with defensins, where genital types differ in their sensitivities to HD5 and HNP1 and where HD5 inhibits multiple steps of HPV16 entry [40-42].

By blocking the externalization of VP1u, the PLA_2_ domain is unable to access the limiting membrane, likely trapping the virus within the endosome. This interpretation is based on the failure of defensin-treated virus to reach the nucleus; however, we did not measure endosomal escape directly. It is plausible that defensins bound to the capsid could mediate endosome escape through their ability to form pores in lipid bilayers [89]. In this case the absence of externalized nuclear localization signals found in VP1u and the VP1/2 common region would still prevent the virus from reaching and entering the nucleus [52, 62]. Thus, the increased amount of detectable virus in defensin-treated samples at 24 h p.i. could be explained by defensin-induced viral aggregation in either the endosome or cytoplasm, protection of virus trapped in the endolysosomal system from lysosomal degradation, or protection of virus in the cytoplasm from ubiquitination and proteosomal degradation. These mechanisms are not mutually exclusive, although we note that all samples are treated with the proteosome inhibitor doxorubicin, which has been shown to increase AAV transduction efficiency [90].

Without additional experimentation, it is difficult to anticipate the generalizability of our findings to other AAV serotypes. Both AAV2 and AAV6 are neutralized by both HD5 and HNP1 when added to the virus either pre- or post-attachment to the cell [22]. However, the sensitivity of AAV1 to each defensin depends on the order-of-addition, where HD5 is only neutralizing when added pre-attachment and HNP1 is only neutralizing when added post-attachment. As described above, the identity of residue 531 in VP3 solely dictates the defensin sensitivities of AAV1 and AAV6. Although the defensin-dependent phenotypes of AAV2 and AAV6 are more similar, AAV2 and AAV1 (not AAV6) have the same residue (glutamic acid) at this position. Thus, the identity of the residue at position 531 is not predictive. AAV2 and AAV6 share the ability to utilize HSPG as an attachment factor [64], while AAV1 uses sialic acid [65]. Therefore, the nature of the attachment factor might correlate with defensin sensitivity, particularly regarding blocking cell binding. Until more serotypes are studied and a completely defensin-resistant serotype is identified, it is hard to determine which characteristics might foretell defensin sensitivity; however, our findings provide a blueprint for key assays that will inform neutralization mechanisms.

Finally, our findings may inform the development of AAV vectors. HNP levels are increased in patients with highly inflammatory disease states, such as cystic fibrosis, which would impair the delivery of gene therapy by defensin-sensitive serotypes such as AAV2 [37]. By identifying where in the entry pathway innate immune peptides such as defensins inhibit, mutant vectors can be better designed with the goal of escaping neutralization. Alternatively, existing vectors that have been customized for tropism or antibody escape can be further tested for defensin sensitivity [8]. Importantly, our findings provide two points of likely inhibition, cell binding and endosome escape, that can be targeted in the development of escape variants.

## Materials and Methods

### Cells

HeLa (ATCC) and HEK293 cells (ATCC or University of Florida Powell Gene Therapy Center Vector Core) were cultured in Dulbecco’s modified Eagle’s media (DMEM) supplemented with 10% Fetal Bovine Serum (FBS), 0.1 mM non-essential amino acids, 4 mM L-glutamine, 100 units/mL penicillin, and 100 µg/mL streptomycin.

### Vector production and purification

AAV2 and AAV6 vectors expressing GFP were produced by triple transfection in mammalian HEK293 cells. HEK293 cells were cultured to approximately 80% confluency in 150 mm cell culture dishes and triple transfected with an equimolar ratio of pXRAAV2 [91], pTR-UF11-GFP [92], and pXX6 (pHelper) [93] plasmids (totaling 40 µg/plate), which were diluted with 1 mL Optimem (Gibco) and 125 μL of 1 mg/ml polyethyleneimine (PEI) per plate. For AAV6, pXR6 [64] was substituted for pXRAAV2. The solution was incubated for 15 min at RT and added dropwise to each plate. Cells were harvested 72 h post-transfection and separated from the supernatant by centrifugation at 1100 × g in a JA-20 rotor at 4°C for 15 min. The cell pellet was resuspended in 1 mL/plate of TD buffer (1X PBS, 5 mM MgCl_2_, 2.5 mM KCl, pH 7.4) and lysed by three freeze/thaw cycles, followed by Benzonase (Promega) treatment at 37°C for 1 h. The cellular debris was removed by centrifugation at 3000 × g, and the clarified cell lysate was stored at -80°C. The supernatant was PEG precipitated overnight with 10% (w/v) PEG 8000 (Fisher) and then centrifuged at 14,300 × g in a JA-10 rotor at 4°C for 90 min. The resulting PEG pellet was resuspended in 1 mL/plate of TD buffer and clarified by centrifugation at 3000 × g. The supernatant was then combined with the cell lysate, and AAV capsids were purified by AVB affinity chromatography on a 1 mL AVB Sepharose affinity chromatography column (Cytvia, Marlborough, MA), as previously described [64, 94]. Virus was eluted with 10 mL of elution buffer (0.1 M glycine-HCl, pH 2.7) and neutralized with 1 mL of neutralization buffer (1 M Tris-HCl, pH 10.0). The final sample was concentrated and buffer exchanged using an Apollo concentrator (Orbital Bioscience), and the purity was verified by SDS-PAGE.

For confocal microscopy, purified AAV2 at approximately 0.5 mg/mL was extensively dialyzed into PBS and labeled with the DyLight 488 Antibody Labeling Kit (Thermo Fischer) according to the manufacturer’s protocol. The completed reaction was dialyzed three times into PBS. Labeled virus was visualized by SDS-PAGE using UV light to detect the fluorophore and quantified by Coomassie Blue staining against BSA standards.

### α-defensin peptides

HD5 (ATCYCRTHECATRESLSGVCEISGRLYRLCCR) and HNP1 (ACYCRIPACIAGERRYGTCIYQGRLWAFCC) were synthesized by CPC Scientific (Sunnyvale, CA) and processed as previously described [45]. In brief, peptides were folded and stabilized by thiol-disulfide reshuffling and purified by reverse-phase high-pressure liquid chromatography (RP-HPLC) to homogeneity. Purity was analyzed by analytical RP-HPLC, verified by mass spectrometry, and quantified by UV absorbance at 280 nm. HD5abu synthesis, purification, and validation have been described previously [20, 21].

### Primary antibodies

A1 (American Research Products, 03-61056) binds an epitope in VP1u [54, 55]. A20 (American Research Products, 03-61055) binds an epitope in the VP1/VP2/VP3 common domain of intact AAV2 [55]. ADK1a (University of Florida Hybridoma Core) binds an epitope in the VP1/VP2/VP3 common domain of intact AAV1 and AAV6 [57, 95]. B1 (University of Florida Hybridoma Core) binds an epitope in the VP1/VP2/VP3 common domain of uncoated AAV2 and AAV6 [54, 55].

### Infection assays

Infection assays were performed as previously described [22]. All viruses and defensins were diluted in serum-free DMEM (SFM) supplemented with 1 µM doxorubicin (Dox). AAV2 was diluted to 5.4E+10 viral genomes (vg)/mL for pre-attachment and 1.4E+11 vg/mL for post-attachment. In brief, for pre-attachment exposure of AAV2 to defensins, 28 µL of AAV2 and 28 µL of SFM with or without HD5 or HNP1 were mixed and incubated on ice for 45 min before 50 µL of the mixture was added to cells. Samples were then incubated at 37°C. For post-attachment exposure of AAV2 to defensins, 50 µL AAV2 was added to chilled cells and allowed to bind for 1 h at 4°C. Samples were washed twice with ice-cold SFM before 50 µL SFM with or without HD5 or HNP1 was added. Defensins were bound at 4°C for 45 min before cells were moved to 37°C. For both conditions, 10 µL 60% FBS in SFM + 1 µM Dox was added to the wells 6 h p.i., but no additional washes were performed. At 48 h p.i., cells were washed with phosphate-buffered saline (PBS), and total monolayer GFP fluorescence was quantified with an Azure Sapphire. Background-subtracted total monolayer fluorescence was further analyzed with Fiji (version 2.14.0/1.54f). Data are shown as the percentage of control infection in the absence of defensin.

### SPR

SPR experiments were performed on a BIAcore X100 instrument at 25°C, with PBS as the running buffer, as previously described [22]. AAV2 was coupled to the CM5 sensor chip using amine coupling, and 9451 response units (RUs) were immobilized. HD5 or HNP1 was the analyte, and the same sensor chip was used for 2 replicate analyses on different days. Data were analyzed as described previously to determine *K*_D_, using the 1:1 binding model in BIAcore X100 evaluation software (version 2.0.0), and stoichiometry, using GraphPad Prism (version 10.3.0).

### Cell binding assays

HeLa cells were detached using 0.5 M EDTA, washed twice with cold SFM, and resuspended in SFM + 0.2% sodium azide (SA). 8.4E+9 particles of AAV2 or AAV6 were incubated with SFM + 0.2% SA with or without twice the final concentration of HD5, HNP1, or ADK1a in a volume of 56 µL. Following incubation for 45 min on ice, 50 µL of the virus/defensin mixture was added to 5.0E+5 HeLa cells in a final volume of 100 µL. For studies with wheat germ agglutinin (WGA), cells were incubated with 50 µg/mL WGA for 10 min on ice, washed, and then incubated with virus plus 50 µg/mL WGA. Samples were incubated for 90 min on ice with intermittent agitation to allow binding. One sample without defensin was reserved as an input control for quantitation. Remaining samples were washed three times with ice cold DMEM. All samples were lysed and processed to extract genomic DNA using the Genejet Genomic DNA Purification kit (Thermofisher) per the manufacturer’s instructions. DNA was eluted from the column in 100 µL elution buffer. qPCR was performed using SsoAdvanced SYBR Green master mix (Bio-Rad) with primers for GFP (F: 5’-CCCAGACCATATGAAGCAGC-3’; R: 5’-GCCCTTAGCTCGATTCTCT-3’). The absolute number of genomes bound to cells in each condition was determined relative to a standard curve produced by a dilution series of the input control. Note that diluting experimental samples 10-fold prior to analysis was required for accurate quantitation. The amount of virus bound to cells in each condition within each replicate was then normalized to the amount of virus bound to control cells in the absence of defensin.

### Immunoprecipitation and western blot

HeLa cells were seeded at 6.0E+5 cells/mL in 6-well plates and cultured overnight. Cells were infected with AAV2 or AAV6 at an MOI of 65,000 vg/cell in 1 mL cold SFM. For AAV2, one well of cells was infected for each condition (ND, HD5, HNP1); for AAV6, three wells were infected for each condition. After incubation for 1 h at 4°C while rocking, the inoculum was removed and replaced with 1 mL SFM with or without 25 µM HD5, 25 µM HD5abu, or 30 µM HNP1. After an additional incubation for 45 min at 4°C while rocking, plates were shifted to 37°C for 5 h (AAV6) or 6 h (AAV2). Cells were washed two times with ice cold PBS, scaped to detach them from the plate, and concentrated in chilled 1.5 mL Eppendorf tubes by centrifugation at 400 × g for 5 min at 4°C. The pellet was lysed in 100 µL immunoprecipitation (IP) buffer (20 mM Tris, 137 mM NaCl, 1% Triton X-100, 2 mM EDTA, 1X HALT protease inhibitor, pH 8), freeze-thawed three times, and centrifuged at 25,000 × g for 10 min at 4°C. Anti-AAV VP1 (A1, 200 ng) was added to the clarified lysate, and samples were incubated overnight at 4°C with constant agitation. For control samples where defensin was added post-lysis, either HD5 (25 µM) or HNP1 (30 µM) was added with A1. A1-bound AAV2 or AAV6 was precipitated with Protein G-PLUS agarose beads (Santa Cruz Biotech). Anti-AAV2 VP1/2/3 (A20, 200 ng) or anti-AAV6 VP1/2/3 (ADK1a, 300 ng) was then added to the supernatant. Samples were incubated for ≥ 2 h at 4°C with constant agitation and precipitated with Protein G-PLUS agarose beads. For both precipitations, beads were washed three times with IP buffer, and samples were eluted with SDS-loading buffer (3.2% SDS, 100 mM Tris, 0.04% bromophenol blue, 16% glycerol (w/v), 40 mM dithiothreitol, pH 6.8). Heat-denatured (95°C for 5 min) samples were analyzed via SDS-PAGE on 7.5% or 10% polyacrylamide gels, transferred to 0.45 µm nitrocellulose membranes, blocked for 1 h at RT in blocking buffer [3% bovine serum albumin (BSA) and 1% Tween-20 in Tris-buffered saline (TBS): 20 mM Tris, 150 mM NaCl, pH 8.0]. Membranes were incubated with anti-AAV B1 antibody (333 ng/mL) in blocking buffer at 4°C overnight, washed three times in TBS + 1% Tween-20, probed with goat anti-mouse Alexa Fluor 488-conjugated secondary antibody (ThermoFisher, A-11001, 1:1000) in blocking buffer, washed twice with TBS + 1% Tween-20 and once with TBS, and imaged on an Azure Sapphire imager. Bands were quantified using AzureSpot Pro (v2.1.097). For AAV2, data is the ratio of the background subtracted volume of the A1 immunoprecipitated VP3 band divided by the A20 immunoprecipitated VP3 band for each sample. For AAV6, the analogous A1/ADK1a ratio was normalized to the value for the no defensin control.

### Differential Scanning Fluorimetry

The stability of AAV2 was determined by differential scanning fluorimetry (DSF) analysis using a Bio-Rad MyiQ2 Thermocycler to monitor the fluorescence produced by the binding of SYPRO Orange dye (Invitrogen) to hydrophobic regions of the AAV2 capsid upon heat denaturation. The temperature of the assay ranged from 30°C to 99°C, ramping at 0.5°C per step. Purified AAV2 (1 µg) was diluted in 22.5 µL of either 25 µM HD5 or 30 µM HNP1 in citrate-phosphate (0.1 M citric acid, 0.2 M Na_2_HPO_4_) buffers at pH 7.4, pH 6, pH 5.5, or pH 4. Samples were incubated for 30 min at RT before 2.5 µl/sample of 1% SYPRO Orange dye was added. Samples containing buffer alone were used as negative controls. Data is the inverse of the negative rate of change of relative fluorescence units vs temperature (−dRFU/dT). The peak value on the resulting thermogram is the T_m_. All experiments were conducted in triplicate.

### Quantification of subcellular localization

HeLa cells were cultured on glass coverslips overnight. DyLight 488-labelled AAV2 in cold SFM containing 1 µM Dox at an MOI of 20,000 vg/cell was added, and cells were incubated for 1 h in the dark at 4°C. The inoculum was removed and replaced with 50 µL SFM containing 1 µM Dox with or without 25 µM HD5 or 30 µM HNP1. After incubation for 45 min at 4°C, one set of control coverslips was washed with PBS before being fixed in 2% PFA (Electron Microscopy Science, 15710) for 15 min at RT. The remaining cells were shifted to 37°C. At 6 h p.i., coverslips were moved to normal growth media, cultured for an additional 18 h, washed with PBS, and fixed. Fixed cells were washed once with PBS and incubated for 20 min at RT in permeabilization buffer (20 mM glycine, 0.5% Triton X-100, PBS). Cells were sequentially stained with AF647-conjugated phalloidin (Abcam, ab176759, 1:1000) in 1% BSA, 0.05% Tween-80 in PBS for 45 min at RT then with 500 ng/mL DAPI for 5 min. Coverslips were mounted with Prolong Gold (Life Technologies, P36930).

Z-series of images spanning entire cells were obtained using a confocal laser-scanning microscope (Zeiss LSM 800) using a 63× objective. A maximum intensity z-projection of each color channel for each field of view was generated using Fiji (v2.14.0/1.54h). Background thresholds for AAV2 signal were determined manually for each replicate in Fiji using uninfected negative controls. A human-in-the-loop model was trained in CellPose 2.0 [96] to identify cell borders and generate cell masks. CellPose masks and z-projection images were imported into CellProfiler (v4.2.5) [97], and background-subtracted single-cell images were identified and exported. A second CellProfiler pipeline was then used to analyze the single-cell images. Nuclei were identified using DAPI by thresholding using the Otsu method. A tertiary object extending 5 pixels into and 50 pixels beyond the nucleus was then defined as the perinuclear space, and the integrated intensities of DyLight 488 within the whole cell, in the perinuclear space, and in the nucleus were measured for each cell. Whole cells with DyLight 488 intensity < 100 relative fluorescence units (RFU) were excluded from analysis.

### Statistical analysis

All statistical analysis was performed with GraphPad Prism 10.3.0. Fig 1A, 1B, 2A, 2C, 2D, and 4A were analyzed by nonlinear regression using the “log (inhibitor) versus response – variable slope (four parameters)” model with the bottom constrained to “0”. Hill slopes and IC_50_ values are the best-fit values. Log IC_50_ values were compared between protocol conditions and assays (e.g., cell binding and inhibition of infection) using the extra-sum-of-squares F test, and the P values are given. Fig 2B and 4B were analyzed by nonlinear regression using the “log (agonist) versus response – variable slope (four parameters)” model without constraint. Note that the AAV6 infection data in Fig 4A and 4B is reproduced from our prior study for comparison [22]. In Fig 3B, 3K, 4C, 4F, and 4G, repeated measures one-way ANOVA using Dunnett’s multiple comparisons test with a single pooled variance was used to compare the mean of each column with that of the “no defensin” control. In Fig 3D, 3F, and 3H a two-tailed, paired t test was used to compare HD5 or HNP1 samples with “no defensin” controls. In Fig 3K, HD5 and HD5abu were analyzed in parallel, while HNP1 was analyzed in separate experiments with distinct “no defensin” controls. HD5, HD5abu, and HNP1 samples are graphed together for clarity. Fig 5C was analyzed by repeated measures two-way ANOVA using Dunnett’s multiple comparisons test with a single pooled variance to compare the mean of each column with that of the “no defensin” control for each pH value. Fig 6A, 6B, 6E and 6F were analyzed by ordinary one-way ANOVA using Dunnett’s multiple comparisons test with a single pooled variance to compare the mean of each column with that of the “no defensin” control. In Fig 6A, each replicate was analyzed individually. In Fig 6B, 6E, and 6F, the data for each condition for all replicates was pooled prior to analysis. In Fig 6E and 6F, data was log-transformed prior to analysis. For all analyses by ANOVA, P values are adjusted to account for multiple comparisons.

## Acknowledgements

The authors would like to dedicate this study to the memory of Dr. Agbandje-McKenna, who was a pioneer in the field of AAV structural biology. We thank Qun Tang and Melike Caglayan for technical assistance with SPR experiments. We thank Dr. Matthew Parsek for the generous use of his microscope and Dr. Jeffrey Carey and Dr. Xuhui Zheng in the Parsek lab for technical assistance with microscopy. This work was supported by R01 AI104920 (to J.G.S. and R.M.) and F32 AI178920 (to K.R.H.), from the National Institute of Allergy and Infectious Diseases (www.niaid.nih.gov) and by the Office of the Director, National Institutes of Health (www.nih.gov/institutes-nih/nih-office-director) under Award Number S10 OD026741 (to J.G.S.). Additional support to J.G.S. in the form of subsidized core services was provided by NIH grants UL1 TR000423 from the National Center for Advancing Translational Sciences (ncats.nih.gov), and P30 CA015704 from the National Cancer Institute (www.cancer.gov). The funders had no role in study design, data collection and analysis, decision to publish, or preparation of the manuscript.

## References

1. Meyer NL, Chapman MS. Adeno-associated virus (AAV) cell entry: structural insights. Trends Microbiol. 2022;30(5):432–51. Epub 20211025. doi: 10.1016/j.tim.2021.09.005. PubMed PMID: 34711462.

2. Worner TP, Bennett A, Habka S, Snijder J, Friese O, Powers T, et al. Adeno-associated virus capsid assembly is divergent and stochastic. Nature communications. 2021;12(1):1642. Epub 20210312. doi: 10.1038/s41467-021-21935-5. PubMed PMID: 33712599; PubMed Central PMCID: PMCPMC7955066.

3. Gao G, Vandenberghe LH, Alvira MR, Lu Y, Calcedo R, Zhou X, et al. Clades of Adeno-associated viruses are widely disseminated in human tissues. J Virol. 2004;78(12):6381–8. Epub 2004/05/28. doi: 10.1128/JVI.78.12.6381-6388.2004. PubMed PMID: 15163731; PubMed Central PMCID: PMCPMC416542.

4. Schmidt M, Grot E, Cervenka P, Wainer S, Buck C, Chiorini JA. Identification and characterization of novel adeno-associated virus isolates in ATCC virus stocks. J Virol. 2006;80(10):5082–5. doi: 10.1128/JVI.80.10.5082-5085.2006. PubMed PMID: 16641301; PubMed Central PMCID: PMCPMC1472088.

5. Bossis I, Chiorini JA. Cloning of an avian adeno-associated virus (AAAV) and generation of recombinant AAAV particles. J Virol. 2003;77(12):6799–810. doi: 10.1128/jvi.77.12.6799-6810.2003. PubMed PMID: 12768000; PubMed Central PMCID: PMCPMC156192.

6. Large EE, Silveria MA, Zane GM, Weerakoon O, Chapman MS. Adeno-Associated Virus (AAV) Gene Delivery: Dissecting Molecular Interactions upon Cell Entry. Viruses. 2021;13(7). Epub 20210710. doi: 10.3390/v13071336. PubMed PMID: 34372542; PubMed Central PMCID: PMCPMC8310307.

7. Pillay S, Carette JE. Host determinants of adeno-associated viral vector entry. Curr Opin Virol. 2017;24:124–31. Epub 20170630. doi: 10.1016/j.coviro.2017.06.003. PubMed PMID: 28672171; PubMed Central PMCID: PMCPMC5549665.

8. Lopez-Gordo E, Chamberlain K, Riyad JM, Kohlbrenner E, Weber T. Natural Adeno-Associated Virus Serotypes and Engineered Adeno-Associated Virus Capsid Variants: Tropism Differences and Mechanistic Insights. Viruses. 2024;16(3). Epub 20240312. doi: 10.3390/v16030442. PubMed PMID: 38543807; PubMed Central PMCID: PMCPMC10975205.

9. Muhuri M, Maeda Y, Ma H, Ram S, Fitzgerald KA, Tai PW, et al. Overcoming innate immune barriers that impede AAV gene therapy vectors. J Clin Invest. 2021;131(1). doi: 10.1172/JCI143780. PubMed PMID: 33393506; PubMed Central PMCID: PMCPMC7773343.

10. Rabinowitz J, Chan YK, Samulski RJ. Adeno-associated Virus (AAV) versus Immune Response. Viruses. 2019;11(2). Epub 20190125. doi: 10.3390/v11020102. PubMed PMID: 30691064; PubMed Central PMCID: PMCPMC6409805.

11. Scallan CD, Jiang H, Liu T, Patarroyo-White S, Sommer JM, Zhou S, et al. Human immunoglobulin inhibits liver transduction by AAV vectors at low AAV2 neutralizing titers in SCID mice. Blood. 2006;107(5):1810–7. Epub 20051025. doi: 10.1182/blood-2005-08-3229. PubMed PMID: 16249376.

12. Boutin S, Monteilhet V, Veron P, Leborgne C, Benveniste O, Montus MF, et al. Prevalence of serum IgG and neutralizing factors against adeno-associated virus (AAV) types 1, 2, 5, 6, 8, and 9 in the healthy population: implications for gene therapy using AAV vectors. Hum Gene Ther. 2010;21(6):704-12. doi: 10.1089/hum.2009.182. PubMed PMID: 20095819.

13. Zaiss AK, Cotter MJ, White LR, Clark SA, Wong NC, Holers VM, et al. Complement is an essential component of the immune response to adeno-associated virus vectors. J Virol. 2008;82(6):2727–40. Epub 20080116. doi: 10.1128/JVI.01990-07. PubMed PMID: 18199646; PubMed Central PMCID: PMCPMC2259003.

14. Arjomandnejad M, Sylvia K, Blackwood M, Nixon T, Tang Q, Muhuri M, et al. Modulating immune responses to AAV by expanded polyclonal T-regs and capsid specific chimeric antigen receptor T-regulatory cells. Mol Ther Methods Clin Dev. 2021;23:490–506. Epub 20211028. doi: 10.1016/j.omtm.2021.10.010. PubMed PMID: 34853797; PubMed Central PMCID: PMCPMC8605179.

15. Basner-Tschakarjan E, Mingozzi F. Cell-Mediated Immunity to AAV Vectors, Evolving Concepts and Potential Solutions. Front Immunol. 2014;5:350. Epub 20140723. doi: 10.3389/fimmu.2014.00350. PubMed PMID: 25101090; PubMed Central PMCID: PMCPMC4107954.

16. Lehrer RI, Lu W. alpha-Defensins in human innate immunity. Immunol Rev. 2012;245(1):84–112. Epub 2011/12/16. doi: 10.1111/j.1600-065X.2011.01082.x. PubMed PMID: 22168415.

17. Selsted ME, Ouellette AJ. Mammalian defensins in the antimicrobial immune response. Nat Immunol. 2005;6(6):551–7. Epub 2005/05/24. doi: 10.1038/ni1206. PubMed PMID: 15908936.

18. Wilson SS, Wiens ME, Smith JG. Antiviral mechanisms of human defensins. J Mol Biol. 2013;425(24):4965–80. Epub 20131002. doi: 10.1016/j.jmb.2013.09.038. PubMed PMID: 24095897; PubMed Central PMCID: PMCPMC3842434.

19. Holly MK, Diaz K, Smith JG. Defensins in Viral Infection and Pathogenesis. Annual review of virology. 2017;4(1):369–91. Epub 20170717. doi: 10.1146/annurev-virology-101416-041734. PubMed PMID: 28715972.

20. Rajabi M, Ericksen B, Wu X, de Leeuw E, Zhao L, Pazgier M, et al. Functional determinants of human enteric alpha-defensin HD5: crucial role for hydrophobicity at dimer interface. J Biol Chem. 2012;287(26):21615–27. Epub 20120509. doi: 10.1074/jbc.M112.367995. PubMed PMID: 22573326; PubMed Central PMCID: PMCPMC3381126.

21. Gounder AP, Wiens ME, Wilson SS, Lu W, Smith JG. Critical determinants of human alpha-defensin 5 activity against non-enveloped viruses. J Biol Chem. 2012;287(29):24554–62. Epub 20120525. doi: 10.1074/jbc.M112.354068. PubMed PMID: 22637473; PubMed Central PMCID: PMCPMC3397880.

22. Porter JM, Oswald MS, Sharma A, Emmanuel S, Kansol A, Bennett A, et al. A Single Surface-Exposed Amino Acid Determines Differential Neutralization of AAV1 and AAV6 by Human Alpha-Defensins. J Virol. 2023;97(3):e0006023. Epub 20230314. doi: 10.1128/jvi.00060-23. PubMed PMID: 36916912; PubMed Central PMCID: PMCPMC10062168.

23. Xu C, Wang A, Marin M, Honnen W, Ramasamy S, Porter E, et al. Human Defensins Inhibit SARS-CoV-2 Infection by Blocking Viral Entry. Viruses. 2021;13(7). Epub 20210626. doi: 10.3390/v13071246. PubMed PMID: 34206990; PubMed Central PMCID: PMCPMC8310277.

24. Holly MK, Smith JG. Paneth Cells during Viral Infection and Pathogenesis. Viruses. 2018;10(5). Epub 20180426. doi: 10.3390/v10050225. PubMed PMID: 29701691; PubMed Central PMCID: PMCPMC5977218.

25. Spencer JD, Hains DS, Porter E, Bevins CL, DiRosario J, Becknell B, et al. Human alpha defensin 5 expression in the human kidney and urinary tract. PLoS One. 2012;7(2):e31712. Epub 20120216. doi: 10.1371/journal.pone.0031712. PubMed PMID: 22359618; PubMed Central PMCID: PMCPMC3281003.

26. Wehkamp J, Chu H, Shen B, Feathers RW, Kays RJ, Lee SK, et al. Paneth cell antimicrobial peptides: topographical distribution and quantification in human gastrointestinal tissues. FEBS Lett. 2006;580(22):5344–50. Epub 2006/09/23. doi: 10.1016/j.febslet.2006.08.083. PubMed PMID: 16989824.

27. Ayabe T, Satchell DP, Wilson CL, Parks WC, Selsted ME, Ouellette AJ. Secretion of microbicidal alpha-defensins by intestinal Paneth cells in response to bacteria. Nat Immunol. 2000;1(2):113–8. Epub 2001/03/15. doi: 10.1038/77783. PubMed PMID: 11248802.

28. Brinkmann V, Reichard U, Goosmann C, Fauler B, Uhlemann Y, Weiss DS, et al. Neutrophil extracellular traps kill bacteria. Science. 2004;303(5663):1532-5. doi: 10.1126/science.1092385. PubMed PMID: 15001782.

29. Melle C, Ernst G, Schimmel B, Bleul A, Thieme H, Kaufmann R, et al. Discovery and identification of alpha-defensins as low abundant, tumor-derived serum markers in colorectal cancer. Gastroenterology. 2005;129(1):66–73. doi: 10.1053/j.gastro.2005.05.014. PubMed PMID: 16012935.

30. Müller CA, Markovic-Lipkovski J, Klatt T, Gamper J, Schwarz G, Beck H, et al. Human α-defensins HNPs-1,-2, and-3 in renal cell carcinoma: Influences on tumor cell proliferation. The American journal of pathology. 2002;160(4):1311-24.

31. Huang C, Lou C, Zheng X, Pang L, Wang G, Zhu M, et al. Plasma human neutrophil peptides as biomarkers of disease severity and mortality in patients with decompensated cirrhosis. Liver Int. 2023;43(5):1096–106. Epub 20230128. doi: 10.1111/liv.15520. PubMed PMID: 36648384.

32. Rycyk-Bojarzynska A, Kasztelan-Szczerbinska B, Cichoz-Lach H, Surdacka A, Rolinski J. Human Neutrophil Alpha-Defensins Promote NETosis and Liver Injury in Alcohol-Related Liver Cirrhosis: Potential Therapeutic Agents. J Clin Med. 2024;13(5). Epub 20240222. doi: 10.3390/jcm13051237. PubMed PMID: 38592082; PubMed Central PMCID: PMCPMC10931661.

33. Xu D, Lu W. Defensins: A Double-Edged Sword in Host Immunity. Front Immunol. 2020;11:764. Epub 2020/05/28. doi: 10.3389/fimmu.2020.00764. PubMed PMID: 32457744; PubMed Central PMCID: PMCPMC7224315.

34. Panyutich AV, Panyutich EA, Krapivin VA, Baturevich EA, Ganz T. Plasma defensin concentrations are elevated in patients with septicemia or bacterial meningitis. The Journal of laboratory and clinical medicine. 1993;122(2):202–7. PubMed PMID: 8340706.

35. Hida RY, Ohashi Y, Takano Y, Dogru M, Goto E, Fujishima H, et al. Elevated levels of human alpha -defensin in tears of patients with allergic conjunctival disease complicated by corneal lesions: detection by SELDI ProteinChip system and quantification. Curr Eye Res. 2005;30(9):723–30. doi: 10.1080/02713680591005986. PubMed PMID: 16123017.

36. Gokcinar NB, Karabulut AA, Onaran Z, Yumusak E, Budak Yildiran FA. Elevated Tear Human Neutrophil Peptides 1-3, Human Beta Defensin-2 Levels and Conjunctival Cathelicidin LL-37 Gene Expression in Ocular Rosacea. Ocular immunology and inflammation. 2019;27(7):1174-83. Epub 20180824. doi: 10.1080/09273948.2018.1504971. PubMed PMID: 30142005.

37. Virella-Lowell I, Poirier A, Chesnut KA, Brantly M, Flotte TR. Inhibition of recombinant adeno-associated virus (rAAV) transduction by bronchial secretions from cystic fibrosis patients. Gene Ther. 2000;7(20):1783–9. doi: 10.1038/sj.gt.3301268. PubMed PMID: 11083501.

38. Smith JG, Nemerow GR. Mechanism of adenovirus neutralization by Human alpha-defensins. Cell host & microbe. 2008;3(1):11–9. Epub 2008/01/15. doi: 10.1016/j.chom.2007.12.001. PubMed PMID: 18191790.

39. Smith JG, Silvestry M, Lindert S, Lu W, Nemerow GR, Stewart PL. Insight into the mechanisms of adenovirus capsid disassembly from studies of defensin neutralization. PLoS Pathog. 2010;6(6):e1000959. Epub 20100624. doi: 10.1371/journal.ppat.1000959. PubMed PMID: 20585634; PubMed Central PMCID: PMCPMC2891831.

40. Wiens ME, Smith JG. Alpha-defensin HD5 inhibits furin cleavage of human papillomavirus 16 L2 to block infection. J Virol. 2015;89(5):2866–74. Epub 20141224. doi: 10.1128/JVI.02901-14. PubMed PMID: 25540379; PubMed Central PMCID: PMCPMC4325740.

41. Wiens ME, Smith JG. alpha-Defensin HD5 Inhibits Human Papillomavirus 16 Infection via Capsid Stabilization and Redirection to the Lysosome. mBio. 2017;8(1). Epub 20170124. doi: 10.1128/mBio.02304-16. PubMed PMID: 28119475; PubMed Central PMCID: PMCPMC5263252.

42. Buck CB, Day PM, Thompson CD, Lubkowski J, Lu W, Lowy DR, et al. Human alpha-defensins block papillomavirus infection. Proc Natl Acad Sci U S A. 2006;103(5):1516–21. Epub 20060123. doi: 10.1073/pnas.0508033103. PubMed PMID: 16432216; PubMed Central PMCID: PMCPMC1360544.

43. Dugan AS, Maginnis MS, Jordan JA, Gasparovic ML, Manley K, Page R, et al. Human alpha-defensins inhibit BK virus infection by aggregating virions and blocking binding to host cells. J Biol Chem. 2008;283(45):31125–32. Epub 20080909. doi: 10.1074/jbc.M805902200. PubMed PMID: 18782756; PubMed Central PMCID: PMCPMC2576552.

44. Zins SR, Nelson CD, Maginnis MS, Banerjee R, O’Hara BA, Atwood WJ. The human alpha defensin HD5 neutralizes JC polyomavirus infection by reducing endoplasmic reticulum traffic and stabilizing the viral capsid. J Virol. 2014;88(2):948–60. Epub 20131106. doi: 10.1128/JVI.02766-13. PubMed PMID: 24198413; PubMed Central PMCID: PMCPMC3911681.

45. Hu CT, Diaz K, Yang LC, Sharma A, Greenberg HB, Smith JG. Corrected and republished from: “VP4 Is a Determinant of Alpha-Defensin Modulation of Rotaviral Infection”. J Virol. 2023;97(10):e0096223. Epub 20231003. doi: 10.1128/jvi.00962-23. PubMed PMID: 37787534; PubMed Central PMCID: PMCPMC10617384.

46. Diaz K, Hu CT, Sul Y, Bromme BA, Myers ND, Skorohodova KV, et al. Defensin-driven viral evolution. PLoS Pathog. 2020;16(11):e1009018. Epub 20201124. doi: 10.1371/journal.ppat.1009018. PubMed PMID: 33232373; PubMed Central PMCID: PMCPMC7723274.

47. Nguyen EK, Nemerow GR, Smith JG. Direct evidence from single-cell analysis that human alpha-defensins block adenovirus uncoating to neutralize infection. J Virol. 2010;84(8):4041–9. Epub 20100203. doi: 10.1128/JVI.02471-09. PubMed PMID: 20130047; PubMed Central PMCID: PMCPMC2849482.

48. Tenge VR, Gounder AP, Wiens ME, Lu W, Smith JG. Delineation of interfaces on human alpha-defensins critical for human adenovirus and human papillomavirus inhibition. PLoS Pathog. 2014;10(9):e1004360. Epub 20140904. doi: 10.1371/journal.ppat.1004360. PubMed PMID: 25188351; PubMed Central PMCID: PMCPMC4154873.

49. Bartlett JS, Wilcher R, Samulski RJ. Infectious entry pathway of adeno-associated virus and adeno-associated virus vectors. J Virol. 2000;74(6):2777–85. PubMed PMID: 10684294.

50. Girod A, Wobus CE, Zadori Z, Ried M, Leike K, Tijssen P, et al. The VP1 capsid protein of adeno-associated virus type 2 is carrying a phospholipase A2 domain required for virus infectivity. J Gen Virol. 2002;83(Pt 5):973–8. Epub 2002/04/19. doi: 10.1099/0022-1317-83-5-973. PubMed PMID: 11961250.

51. Kronenberg S, Bottcher B, von der Lieth CW, Bleker S, Kleinschmidt JA. A conformational change in the adeno-associated virus type 2 capsid leads to the exposure of hidden VP1 N termini. J Virol. 2005;79(9):5296–303. Epub 2005/04/14. doi: 10.1128/JVI.79.9.5296-5303.2005. PubMed PMID: 15827144; PubMed Central PMCID: PMCPMC1082756.

52. Sonntag F, Bleker S, Leuchs B, Fischer R, Kleinschmidt JA. Adeno-associated virus type 2 capsids with externalized VP1/VP2 trafficking domains are generated prior to passage through the cytoplasm and are maintained until uncoating occurs in the nucleus. J Virol. 2006;80(22):11040–54. Epub 20060906. doi: 10.1128/JVI.01056-06. PubMed PMID: 16956943; PubMed Central PMCID: PMCPMC1642181.

53. Stahnke S, Lux K, Uhrig S, Kreppel F, Hosel M, Coutelle O, et al. Intrinsic phospholipase A2 activity of adeno-associated virus is involved in endosomal escape of incoming particles. Virology. 2011;409(1):77–83. Epub 20101025. doi: 10.1016/j.virol.2010.09.025. PubMed PMID: 20974479.

54. Wobus CE, Hugle-Dorr B, Girod A, Petersen G, Hallek M, Kleinschmidt JA. Monoclonal antibodies against the adeno-associated virus type 2 (AAV-2) capsid: epitope mapping and identification of capsid domains involved in AAV-2-cell interaction and neutralization of AAV-2 infection. J Virol. 2000;74(19):9281–93. Epub 2000/09/12. doi: 10.1128/jvi.74.19.9281-9293.2000. PubMed PMID: 10982375; PubMed Central PMCID: PMCPMC102127.

55. Wistuba A, Kern A, Weger S, Grimm D, Kleinschmidt JA. Subcellular compartmentalization of adeno-associated virus type 2 assembly. J Virol. 1997;71(2):1341–52. doi: 10.1128/JVI.71.2.1341-1352.1997. PubMed PMID: 8995658; PubMed Central PMCID: PMCPMC191189.

56. Huang LY, Patel A, Ng R, Miller EB, Halder S, McKenna R, et al. Characterization of the Adeno-Associated Virus 1 and 6 Sialic Acid Binding Site. J Virol. 2016;90(11):5219–30. Epub 20160512. doi: 10.1128/JVI.00161-16. PubMed PMID: 26962225; PubMed Central PMCID: PMCPMC4934739.

57. Kuck D, Kern A, Kleinschmidt JA. Development of AAV serotype-specific ELISAs using novel monoclonal antibodies. J Virol Methods. 2007;140(1-2):17–24. Epub 20061128. doi: 10.1016/j.jviromet.2006.10.005. PubMed PMID: 17126418.

58. Li W, Zhang L, Wu Z, Pickles RJ, Samulski RJ. AAV-6 mediated efficient transduction of mouse lower airways. Virology. 2011;417(2):327–33. Epub 20110714. doi: 10.1016/j.virol.2011.06.009. PubMed PMID: 21752418; PubMed Central PMCID: PMCPMC3163804.

59. Salganik M, Venkatakrishnan B, Bennett A, Lins B, Yarbrough J, Muzyczka N, et al. Evidence for pH-dependent protease activity in the adeno-associated virus capsid. J Virol. 2012;86(21):11877–85. Epub 20120822. doi: 10.1128/JVI.01717-12. PubMed PMID: 22915820; PubMed Central PMCID: PMCPMC3486322.

60. Rayaprolu V, Kruse S, Kant R, Venkatakrishnan B, Movahed N, Brooke D, et al. Comparative analysis of adeno-associated virus capsid stability and dynamics. J Virol. 2013;87(24):13150–60. Epub 2013/09/27. doi: 10.1128/JVI.01415-13. PubMed PMID: 24067976; PubMed Central PMCID: PMCPMC3838259.

61. Lins-Austin B, Patel S, Mietzsch M, Brooke D, Bennett A, Venkatakrishnan B, et al. Adeno-Associated Virus (AAV) Capsid Stability and Liposome Remodeling During Endo/Lysosomal pH Trafficking. Viruses. 2020;12(6). Epub 2020/06/25. doi: 10.3390/v12060668. PubMed PMID: 32575696; PubMed Central PMCID: PMCPMC7354436.

62. Grieger JC, Snowdy S, Samulski RJ. Separate basic region motifs within the adeno-associated virus capsid proteins are essential for infectivity and assembly. J Virol. 2006;80(11):5199–210. Epub 2006/05/16. doi: 10.1128/JVI.02723-05. PubMed PMID: 16699000; PubMed Central PMCID: PMCPMC1472161.

63. Summerford C, Samulski RJ. Membrane-associated heparan sulfate proteoglycan is a receptor for adeno-associated virus type 2 virions. J Virol. 1998;72(2):1438–45. Epub 1998/01/28. PubMed PMID: 9445046; PubMed Central PMCID: PMCPMC124624.

64. Wu Z, Asokan A, Grieger JC, Govindasamy L, Agbandje-McKenna M, Samulski RJ. Single amino acid changes can influence titer, heparin binding, and tissue tropism in different adeno-associated virus serotypes. J Virol. 2006;80(22):11393–7. Epub 20060830. doi: 10.1128/JVI.01288-06. PubMed PMID: 16943302; PubMed Central PMCID: PMCPMC1642158.

65. Wu Z, Miller E, Agbandje-McKenna M, Samulski RJ. Alpha2,3 and alpha2,6 N-linked sialic acids facilitate efficient binding and transduction by adeno-associated virus types 1 and 6. J Virol. 2006;80(18):9093–103. Epub 2006/08/31. doi: 10.1128/JVI.00895-06. PubMed PMID: 16940521; PubMed Central PMCID: PMCPMC1563919.

66. Zengel J, Carette JE. Structural and cellular biology of adeno-associated virus attachment and entry. Adv Virus Res. 2020;106:39–84. Epub 20200213. doi: 10.1016/bs.aivir.2020.01.002. PubMed PMID: 32327148.

67. Dudek AM, Pillay S, Puschnik AS, Nagamine CM, Cheng F, Qiu J, et al. An Alternate Route for Adeno-associated Virus (AAV) Entry Independent of AAV Receptor. J Virol. 2018;92(7). Epub 2018/01/19. doi: 10.1128/JVI.02213-17. PubMed PMID: 29343568; PubMed Central PMCID: PMCPMC5972900.

68. Pillay S, Meyer NL, Puschnik AS, Davulcu O, Diep J, Ishikawa Y, et al. An essential receptor for adeno-associated virus infection. Nature. 2016;530(7588):108-12. Epub 20160127. doi: 10.1038/nature16465. PubMed PMID: 26814968; PubMed Central PMCID: PMCPMC4962915.

69. Dudek AM, Zabaleta N, Zinn E, Pillay S, Zengel J, Porter C, et al. GPR108 Is a Highly Conserved AAV Entry Factor. Mol Ther. 2020;28(2):367–81. Epub 20191113. doi: 10.1016/j.ymthe.2019.11.005. PubMed PMID: 31784416; PubMed Central PMCID: PMCPMC7000996.

70. Meisen WH, Nejad ZB, Hardy M, Zhao H, Oliverio O, Wang S, et al. Pooled Screens Identify GPR108 and TM9SF2 as Host Cell Factors Critical for AAV Transduction. Mol Ther Methods Clin Dev. 2020;17:601–11. Epub 20200317. doi: 10.1016/j.omtm.2020.03.012. PubMed PMID: 32280726; PubMed Central PMCID: PMCPMC7139131.

71. Mantyla E, Kann M, Vihinen-Ranta M. Protoparvovirus Knocking at the Nuclear Door. Viruses. 2017;9(10). Epub 20171002. doi: 10.3390/v9100286. PubMed PMID: 28974036; PubMed Central PMCID: PMCPMC5691637.

72. Sanlioglu S, Benson PK, Yang J, Atkinson EM, Reynolds T, Engelhardt JF. Endocytosis and nuclear trafficking of adeno-associated virus type 2 are controlled by rac1 and phosphatidylinositol-3 kinase activation. J Virol. 2000;74(19):9184–96. doi: 10.1128/jvi.74.19.9184-9196.2000. PubMed PMID: 10982365; PubMed Central PMCID: PMCPMC102117.

73. Xiao PJ, Samulski RJ. Cytoplasmic trafficking, endosomal escape, and perinuclear accumulation of adeno-associated virus type 2 particles are facilitated by microtubule network. J Virol. 2012;86(19):10462–73. Epub 20120718. doi: 10.1128/JVI.00935-12. PubMed PMID: 22811523; PubMed Central PMCID: PMCPMC3457265.

74. Sutter SO, Lkharrazi A, Schraner EM, Michaelsen K, Meier AF, Marx J, et al. Adeno-associated virus type 2 (AAV2) uncoating is a stepwise process and is linked to structural reorganization of the nucleolus. PLoS Pathog. 2022;18(7):e1010187. Epub 20220711. doi: 10.1371/journal.ppat.1010187. PubMed PMID: 35816507; PubMed Central PMCID: PMCPMC9302821.

75. Johnson JS, Samulski RJ. Enhancement of adeno-associated virus infection by mobilizing capsids into and out of the nucleolus. J Virol. 2009;83(6):2632–44. Epub 20081224. doi: 10.1128/JVI.02309-08. PubMed PMID: 19109385; PubMed Central PMCID: PMCPMC2648275.

76. Venkatakrishnan B, Yarbrough J, Domsic J, Bennett A, Bothner B, Kozyreva OG, et al. Structure and dynamics of adeno-associated virus serotype 1 VP1-unique N-terminal domain and its role in capsid trafficking. J Virol. 2013;87(9):4974–84. Epub 20130220. doi: 10.1128/JVI.02524-12. PubMed PMID: 23427155; PubMed Central PMCID: PMCPMC3624325.

77. Bleker S, Sonntag F, Kleinschmidt JA. Mutational analysis of narrow pores at the fivefold symmetry axes of adeno-associated virus type 2 capsids reveals a dual role in genome packaging and activation of phospholipase A2 activity. J Virol. 2005;79(4):2528–40. doi: 10.1128/JVI.79.4.2528-2540.2005. PubMed PMID: 15681453; PubMed Central PMCID: PMCPMC546590.

78. Horowitz ED, Finn MG, Asokan A. Tyrosine cross-linking reveals interfacial dynamics in adeno-associated viral capsids during infection. ACS Chem Biol. 2012;7(6):1059–66. Epub 20120406. doi: 10.1021/cb3000265. PubMed PMID: 22458529; PubMed Central PMCID: PMCPMC3376196.

79. Emmanuel SN, Smith JK, Hsi J, Tseng YS, Kaplan M, Mietzsch M, et al. Structurally Mapping Antigenic Epitopes of Adeno-associated Virus 9: Development of Antibody Escape Variants. J Virol. 2022;96(3):e0125121. Epub 20211110. doi: 10.1128/JVI.01251-21. PubMed PMID: 34757842; PubMed Central PMCID: PMCPMC8827038.

80. Havlik LP, Simon KE, Smith JK, Klinc KA, Tse LV, Oh DK, et al. Coevolution of Adeno-associated Virus Capsid Antigenicity and Tropism through a Structure-Guided Approach. J Virol. 2020;94(19). Epub 20200915. doi: 10.1128/JVI.00976-20. PubMed PMID: 32669336; PubMed Central PMCID: PMCPMC7495376.

81. Mietzsch M, Smith JK, Yu JC, Banala V, Emmanuel SN, Jose A, et al. Characterization of AAV-Specific Affinity Ligands: Consequences for Vector Purification and Development Strategies. Mol Ther Methods Clin Dev. 2020;19:362–73. Epub 20201004. doi: 10.1016/j.omtm.2020.10.001. PubMed PMID: 33145372; PubMed Central PMCID: PMCPMC7591348.

82. Kurian JJ, Lakshmanan R, Chmely WM, Hull JA, Yu JC, Bennett A, et al. Adeno-Associated Virus VP1u Exhibits Protease Activity. Viruses. 2019;11(5). Epub 20190429. doi: 10.3390/v11050399. PubMed PMID: 31035643; PubMed Central PMCID: PMCPMC6563295.

83. Farr GA, Cotmore SF, Tattersall P. VP2 cleavage and the leucine ring at the base of the fivefold cylinder control pH-dependent externalization of both the VP1 N terminus and the genome of minute virus of mice. J Virol. 2006;80(1):161–71. doi: 10.1128/JVI.80.1.161-171.2006. PubMed PMID: 16352540; PubMed Central PMCID: PMCPMC1317546.

84. Callaway HM, Feng KH, Lee DW, Allison AB, Pinard M, McKenna R, et al. Parvovirus Capsid Structures Required for Infection: Mutations Controlling Receptor Recognition and Protease Cleavages. J Virol. 2017;91(2). Epub 20170103. doi: 10.1128/JVI.01871-16. PubMed PMID: 27847360; PubMed Central PMCID: PMCPMC5215354.

85. Mani B, Baltzer C, Valle N, Almendral JM, Kempf C, Ros C. Low pH-dependent endosomal processing of the incoming parvovirus minute virus of mice virion leads to externalization of the VP1 N-terminal sequence (N-VP1), N-VP2 cleavage, and uncoating of the full-length genome. J Virol. 2006;80(2):1015-24. doi: 10.1128/JVI.80.2.1015-1024.2006. PubMed PMID: 16379002; PubMed Central PMCID: PMCPMC1346861.

86. Zhao C, Porter JM, Burke PC, Arnberg N, Smith JG. Alpha-defensin binding expands human adenovirus tropism. PLoS Pathog. 2024;20(6):e1012317. Epub 20240620. doi: 10.1371/journal.ppat.1012317. PubMed PMID: 38900833; PubMed Central PMCID: PMCPMC11230588.

87. Wilson SS, Bromme BA, Holly MK, Wiens ME, Gounder AP, Sul Y, et al. Alpha-defensin-dependent enhancement of enteric viral infection. PLoS Pathog. 2017;13(6):e1006446. Epub 20170616. doi: 10.1371/journal.ppat.1006446. PubMed PMID: 28622386; PubMed Central PMCID: PMCPMC5489213.

88. Rapista A, Ding J, Benito B, Lo YT, Neiditch MB, Lu W, et al. Human defensins 5 and 6 enhance HIV-1 infectivity through promoting HIV attachment. Retrovirology. 2011;8:45. Epub 20110614. doi: 10.1186/1742-4690-8-45. PubMed PMID: 21672195; PubMed Central PMCID: PMCPMC3146398.

89. Hazlett L, Wu M. Defensins in innate immunity. Cell Tissue Res. 2011;343(1):175–88. Epub 20100821. doi: 10.1007/s00441-010-1022-4. PubMed PMID: 20730446.

90. Yan Z, Zak R, Zhang Y, Ding W, Godwin S, Munson K, et al. Distinct classes of proteasome-modulating agents cooperatively augment recombinant adeno-associated virus type 2 and type 5-mediated transduction from the apical surfaces of human airway epithelia. J Virol. 2004;78(6):2863–74. doi: 10.1128/jvi.78.6.2863-2874.2004. PubMed PMID: 14990705; PubMed Central PMCID: PMCPMC353734.

91. Rabinowitz JE, Rolling F, Li C, Conrath H, Xiao W, Xiao X, et al. Cross-packaging of a single adeno-associated virus (AAV) type 2 vector genome into multiple AAV serotypes enables transduction with broad specificity. J Virol. 2002;76(2):791–801. doi: 10.1128/jvi.76.2.791-801.2002. PubMed PMID: 11752169; PubMed Central PMCID: PMCPMC136844.

92. Burger C, Gorbatyuk OS, Velardo MJ, Peden CS, Williams P, Zolotukhin S, et al. Recombinant AAV viral vectors pseudotyped with viral capsids from serotypes 1, 2, and 5 display differential efficiency and cell tropism after delivery to different regions of the central nervous system. Mol Ther. 2004;10(2):302-17. doi: 10.1016/j.ymthe.2004.05.024. PubMed PMID: 15294177.

93. Xiao X, Li J, Samulski RJ. Production of high-titer recombinant adeno-associated virus vectors in the absence of helper adenovirus. J Virol. 1998;72(3):2224–32. doi: 10.1128/JVI.72.3.2224-2232.1998. PubMed PMID: 9499080; PubMed Central PMCID: PMCPMC109519.

94. Bennett A, Patel S, Mietzsch M, Jose A, Lins-Austin B, Yu JC, et al. Thermal Stability as a Determinant of AAV Serotype Identity. Mol Ther Methods Clin Dev. 2017;6:171–82. Epub 20170724. doi: 10.1016/j.omtm.2017.07.003. PubMed PMID: 28828392; PubMed Central PMCID: PMCPMC5552060.

95. Tseng YS, Gurda BL, Chipman P, McKenna R, Afione S, Chiorini JA, et al. Adeno-associated virus serotype 1 (AAV1)- and AAV5-antibody complex structures reveal evolutionary commonalities in parvovirus antigenic reactivity. J Virol. 2015;89(3):1794–808. Epub 20141119. doi: 10.1128/JVI.02710-14. PubMed PMID: 25410874; PubMed Central PMCID: PMCPMC4300747.

96. Pachitariu M, Stringer C. Cellpose 2.0: how to train your own model. Nat Methods. 2022;19(12):1634–41. Epub 20221107. doi: 10.1038/s41592-022-01663-4. PubMed PMID: 36344832; PubMed Central PMCID: PMCPMC9718665.

97. Stirling DR, Swain-Bowden MJ, Lucas AM, Carpenter AE, Cimini BA, Goodman A. CellProfiler 4: improvements in speed, utility and usability. BMC Bioinformatics. 2021;22(1):433. Epub 20210910. doi: 10.1186/s12859-021-04344-9. PubMed PMID: 34507520; PubMed Central PMCID: PMCPMC8431850.

